# Allelic Association Analyses: Estimation Recommendations

**DOI:** 10.64898/2026.01.26.701864

**Authors:** Bruce S. Weir, Jérôme Goudet

## Abstract

We review the rich literature on the estimation of measures of inbreeding, relatedness and population structure, beginning with Sewall Wright’s *F*-statistics and moving onto the descriptive statistics of Masatoshi Nei and Clark Cockerham. The current availability of genome-level single nucleotide variant data is allowing for sophisticated treatments of inferred identity by descent segments and inferred ancestral recombination graphs. Underlying such disparate methods is an emphasis of characterizing the descent status of alleles within and between individuals and populations and we have found allele-sharing statistics a convenient framework for examining the differences and similarities among different estimators. We have been able to resolve some long-standing reported differences among estimators, especially those involving the work of Nei.

In the course of our algebraic and empirical treatment of descent measure estimation we have been able to formulate a set of five recommendations. Following the early work of Sewall Wright, we recommend *1. State that descent measures for pairs of alleles are relative to values in a reference set of allele pairs*. With this view, we recommend *2. Use estimators that preserve descent measure rankings over different reference sets*. Allele-sharing estimators satisfy this recommendation. Reducing genotypic data to allelic data has the benefit of reducing dimensionality, but we recommend *3. If genotypic data are available, avoid having to assume Hardy-Weinberg equilibrium by not reducing them to allelic data*. Partly as a consequence of working with genotypic data, we recommend *4. Recognize that allele frequencies do not need to be estimated*. Not estimating allele frequencies prevents the confounding of descent estimates for target pairs of alleles by the status of all pairs in a reference set. On the basis of both theoretical and empirical results, finally we recommend *5. Consider both inbreeding and kinship when estimating either one*. It is difficult to envisage a natural population with relatedness but no inbreeding, or vice versa.

## Introduction

There is a rich population and quantitative genetics literature on methods for estimating descent measures: characterizations of inbreeding, relatedness and population structure. Although the single-locus methods of Wright (1922), Malécot (1948), Nei (1973), Weir and Cockerham (1984) (WC84 hereafter), Yang et al. (2011), Weir and Goudet (2017) (WG17 hereafter), Hou and Ochoa (2023) (HO23 hereafter) and others appear quite different at first, they all rest on statements about pairs of alleles within or between individuals, and pairs of alleles within or between populations. Here we present algebraic translations from one set of parameters to another to show that differences are more apparent than real, and often disappear completely for estimates based on large sample sizes within populations or equal sample sizes among populations. Our discussion is especially timely in light of the recent passing of Masatoshi Nei, a major contributor to this field. His work remains relevant fifty years after his early publications. Throughout this paper we lay the foundations for this set of five recommendations:

1. State that descent measures for pairs of alleles are relative to values in a reference set of allele pairs.
2. Use estimators that preserve descent measure rankings over different reference sets.
3. If genotypic data are available, avoid having to assume Hardy-Weinberg equilibrium by not reducing them to allelic data.
4. Recognize that allele frequencies do not need to be estimated.
5. Consider both inbreeding and kinship when estimating either one.

The foundational work in this area was by Sewall Wright. He described his *F*-statistics in his 1951 and 1965 papers (Wright 1951, 1965) as the correlations of pairs of gametes within or between individuals relative to random pairs of alleles within a population or from a set of populations. We keep the style of Wright’s definitions but prefer language that stresses the comparisons of pairs of distinct alleles at different levels in a hierarchy of individuals and populations (Cockerham and Weir 1987; Hudson *et al*. 1992; Rousset 1996):

> *F*_*IT*_ is the correlation between pairs of alleles that unite to produce the individuals, relative to random pairs of alleles, one from each of two distinct populations in a set of populations. *F*_*IS*_ is the average over all populations of the correlation between uniting alleles relative to pairs of alleles, one from each of two distinct individuals in the same population. *F*_*ST*_ is the correlation between pairs of alleles, one from each of distinct individuals within a population, relative to random pairs of alleles, one from each of distinct populations of the set of populations.

These modified correlations reflect the consequences of pairs of alleles being within individuals or being in pairs of distinct individuals within or between populations, and do not depend on how many individuals or populations there are in a study.

We focus on single nucleotide variants and on a designated allele, such as the ancestral allele, for each variant. For variants with more than two alleles we take averages over alleles. This lets us regard the correlations for Wright’s *F*-statistics as being for allelic indicators *x* that have the value 1 if an observed allele is the designated type and are zero otherwise, as stated by Hill (1996). The sum *X* of the two indicators for the pair of alleles carried by a diploid individual is the allele dosage that individual has for the designated allele. We discuss other ploidy levels below. The expected value of *x* over all evolutionary replicates of the history of a variant, from some initial or reference population, is *π*, the probability the variant is the designated type: we find it helpful to use the symbol *π* different from the usual *p* to emphasize the difference between unknown parameters and potentially observable quantities (Zhang *et al*. 2022). The same expectation with respect to a reference population applies to the correlation between pairs of indicator variables. It follows (Cockerham 1973) that the probability a pair of variants have the designated allele type is *π*^2^ + *π*(1 − *π*)*θ* where *θ* is the correlation of the indicator variables for that pair of alleles. Alternatively, following Malécot (1948), we can regard *θ* for a pair of variants as the probability they are identical by descent (ibd) rather than as the correlation of their allelic indicators. Identity by descent also requires a reference population, in which generally all alleles are considered to be not ibd. The correlation and ibd approaches are broadly equivalent, as noted by Wright himself (Wright 1965), although ibd probabilities are necessarily positive. We use the term “descent measure” to refer to either allelic state correlation or allelic probability of identity by descent. Rousset (1996) used an equivalent formulation with probabilities of pairs of alleles being identical in state. Neither correlations nor identity probabilities themselves are estimable from data collected from only a set of contemporary populations, as implied by “There is no absolute measure of ibd: ibd is always relative to some reference population.” (Thompson 2013) and used in our Recommendation 1. The need for a reference population was also discussed by Cotterman (1940) and Rousset (2002).

We refer to several recent publications as appropriate in the text below, but note here that we have found the writings of Wang (2014, 2017, 2022) to be very useful. He discusses the consequences of using sample allele frequencies for descent measure estimation and our own use of allele-sharing statistics is a direct response to his statement in Wang (2017)

> The estimators used for this purpose [relatedness estimation] assume that marker allele probabilities in a population are known without error. Unfortunately, however, these frequencies, upon which both the definition and the estimation of relatedness are based, are rarely known in reality.

We agree with Wang that relatedness is best defined by comparing descent status of target and reference alleles, as we detail below, although we disagree slightly with him by showing that estimators can still be expressed as functions of probability of identity by descent. Rather than trying to estimate both allele probabilities and descent measures with data from extant populations, we avoid the estimation of allele probabilities (Recommendation 4). Most recently, Wang (2025) has used an iterative procedure to estimate both allele probabilities and ibd coefficients, with computational efficiencies for small samples or for samples with high proportions of highly related individuals.

We also note several similarities between the allele-sharing approach and the work of Ritland (1996). His allele-specific proportion of alleles of each type *u* can be combined over types to give the allele-sharing statistics we employ. He uses sample allele frequencies but acknowledges that this does not account for the inbreeding and relatedness among all individuals in the sample. Recently he extended his approach to sets of three or four alleles (Ritland 2024).

## Materials and Methods

### Classes of Allele Pairs

We distinguish among different classes of pairs of alleles with suffices on the *θ*’s, as shown in Box 1. In particular, that box shows 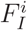 as the probability a random individual in population *i* carries correlated or ibd alleles for any variant and, from the expression in the previous paragraph, the probability 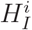 a random individual in that population is heterozygous is

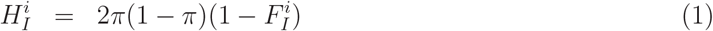

The parameters 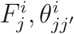 for individuals *j* and individual-pairs *j, j*′, and their averages 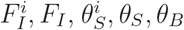 over individuals and populations, are foundational in the sense that they provide values for the other averages 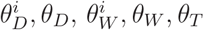 shown in Box 1.

Box 1

We use unweighted averages over populations in Box 1 as we do not expect the *θ* values to depend on sample sizes. We have used sample size weights in the past (WC84) because we assumed at that time the same values of *θ* applied to all populations. We subsequently (Weir and Hill 2002) relaxed that assumption.

To be consistent with Wright we use subscript *I* for alleles within an individual and *T* for random pairs of alleles from the set of populations in a study, but we use *S* for pairs of alleles from distinct individuals and *W* for random pairs of alleles from a single population. Descent measures for alleles from distinct populations are identified by *B*. The measures in Box 1 allow Wright’s *F*-statistics *F* ^W^ to be written as

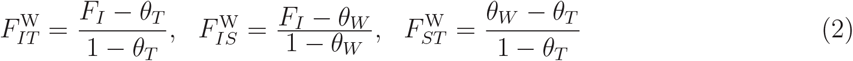

This formulation not only helps explain the meaning of Wright’s “relative to” but also it shows that the inbreeding coefficient 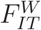 contrasts pairs of alleles within individuals with random pairs of alleles. The evolutionary forces that affect correlations of alleles within individuals are better accommodated, however, by making the correlations relative to alleles from different individuals in the same or different populations to give alternative *F*-statistics that we write as *F* ^AS^ since we estimate them with allele-sharing statistics:

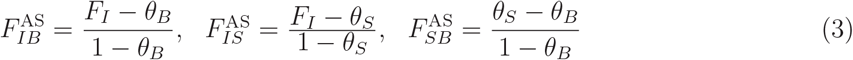

Wright’s relationship 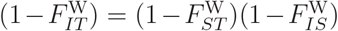 is confirmed by Equations 2, and the analogous result 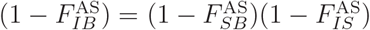 follows from Equations 3.

It is not possible to have a notation that allows the *θ*’s for the *F*-statistics in Equations 2 and 3 to have the same subscripts in both sets, and for those subscripts to match Wright’s *I, S, T* subscripts on the *F* ‘s. Recasting Wright’s widely-used *F*_*ST*_ with the allele-sharing *F*_*SB*_ risks confusion but it does stress our emphasis on comparisons with alleles in distinct individuals or populations and it provides nmemonic value in later estimation discussion.

There may be another nomenclature issue in Equations 2 and 3 since “*F*_*ST*_ “ has often been defined in terms of observable statistics (e.g. Hartl and Clark 2007). We sought previously (WG17; Cockerham and Weir 1987) to make a distinction between statistics and parameters by using the symbol *β* in place of *F* for parametric quantities like (*θ*_*S*_ − *θ*_*B*_)/(1 − *θ*_*B*_) and 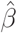 for their estimators. We now suggest distinguishing parameters and their estimators by a caret for estimators, and stress that both parameters and estimators are compound “relative to” quantities. We have previously also used *M* or *Q* (WG17; Cockerham and Weir 1987) for allele-sharing, but here and in Zhang *et al*. (2022) we join Ochoa and Storey (2021) in using *A*. The *Q* notation has also been used by Rousset (2007), who defined a general class of *F*-statistics as (*Q*_*w*_ − *Q*_*b*_)/(1 − *Q*_*b*_) in terms of identity by descent for alleles within (w) and between (b) specified classes of alleles, and estimated these ratios by using allele-sharing statistics for these classes of alleles.

Equations 3 do not make explicit use of sample sizes or numbers of populations, but these dimensions do affect Wright’s coefficients. From Box 1, for sample sizes *n*_*i*_ large enough that 1*/n*_*i*_ terms can be ignored and *θ*_*W*_ = *θ*_*S*_, for example:

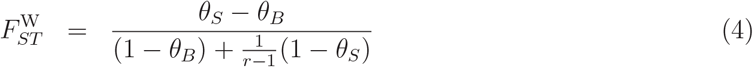

where *r* is the number of populations. Only for large numbers *r* of populations is 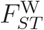 close to 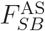, and 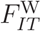 close to 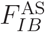, although 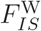 is close to 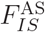 for large sample sizes *n*.

The descent coefficients *θ* can also be defined for specific individuals and specific pairs of individuals within populations. The correlation for alleles uniting to form an individual is the inbreeding coefficient *F* for that individual and may be distinguished according to population with a superscript, and according to individual with a subscript. Various averages may be taken over individuals or populations. The identity for pairs of alleles, one each from two distinct individuals within a population, is the coancestry coefficient for those individuals. Population superscripts and individual-pair subscripts may be appended. The “relative to” nature of inbreeding and coancestry coefficients is made explicit by comparing the relationship measure for a target pair of alleles to the average measure for pairs of alleles in some reference set of alleles. When there is no need to specify that an allele-sharing approach is being used and AS superscripts are not needed, the within-population individual-specific inbreeding coefficients are

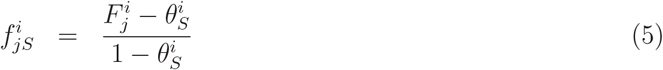

with an average over individuals of 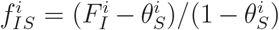. Because the population-specific value 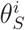 may vary over populations, we would need to weight the 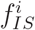 by the corresponding 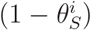 to get an average with value of 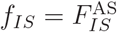 shown in Equation 3.

The within-population individual-pair-specific kinship coefficients are

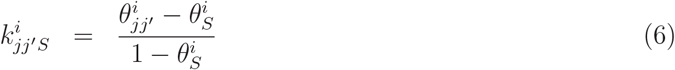

As 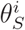 is the average of the 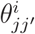 over all pairs of distinct individuals within population *i*, the average within-population kinship is zero, although the average within-population inbreeding coefficient can be positive or negative. The reference set of allele pairs in Equations 5 and 6 can be changed from those within a population to those in the whole study by removing population labels: *θ*_*jj*_′ is then for any pair of individuals *j, j*′ and *θ*_*S*_, now written as *θ*_*S*_∗ to emphasize it refers to the whole study, is the average coancestry for all pairs in the study.

An alternative reference set for pairs of alleles within populations consists of pairs between populations, as is appropriate for the *F*-statistics used to characterize populations. The reference set ibd is then *θ*_*B*_ and the within-population inbreeding and kinships are 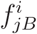 and 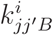 with averages over populations of 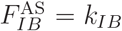 and 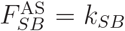 shown in Equations 3 for the set of populations in a study.

The key result is that the descent measures for different pairs of alleles in a set of alleles have the same rankings for all reference sets of allele pairs. The *θ*’s themselves cannot be estimated without knowing the allele probabilities *π* but they can be transformed to equally-ranked values relative to those in a reference set.

### Genetic Relatedness Matrix

Measures of inbreeding and relatedness among sets of individuals are necessary in quantitative genetic analyses of the influence of genetic profiles on trait values. If the trait values for a set of *n* individuals is written in vector form as ***Y***, then the effect of a genetic variant can be described by a regression coefficient in a linear mixed model (Yang *et al*. 2011):

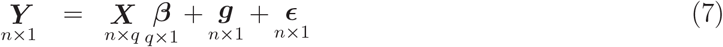

where ***β*** is a set of *q* fixed effects, including a mean, an effect of a particular variant and other effects such as sex and age, ***X*** is a set of coefficients, ***g*** is a set of random effects representing the whole genetic profiles of study individuals and ***ϵ*** is a set of residuals generally considered to be independent across individuals and identically normally distributed. If the genetic model for traits has only single-locus additive effects, the *n × n* variance matrix of the vector of observations is

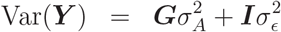

where 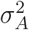 is the (additive component of) genetic variance, 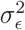 is the residual error and ***I*** is the *n × n* identity matrix. If the *n* observed individuals are from population *i*, the *n × n* genetic relatedness matrix (GRM) ***G*** has elements 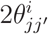 for row *j* and column *j*′. Diagonal elements 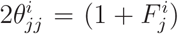 reflect individual inbreeding levels and off-diagonal elements are twice the coancestry coefficients for pairs of individuals. The GRM is also called the numerator relationship matrix (NRM) when the elements are determined from pedigrees and ***K*** = ***G***/2 is called the kinship matrix. A special case of interest is when the trait value for an individual is its allele dosage at a SNP and there is no error term: the additive genetic variance component is then 2*π*(1 − *π*) for that locus.

The linear model parameters of interest are ***β***, 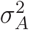 and 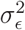. We now consider ways to assign numerical values to ***G*** to allow their estimation: historically these values were predicted from known pedigrees of studied individuals (Yu *et al*. 2006).

### Estimation

#### Sample Allele Frequency Methods

The focus of this paper is on making inferences about allelic descent measures from data, with an emphasis on the relationships among the various approaches. Until recently, the analysis of population structure was framed in terms of sample allele frequencies, as in Wright (1943, 1965), Li and Horvitz (1953), Nei (1973), WC84, VanRaden (2008) and Yang *et al*. (2010). Manichaikul *et al*. (2010) avoided using allele frequencies but assumed Hardy-Weinberg equilibrium, as we discuss below. Another alternative to using allele frequencies is to regard long runs of homozygosity within individuals, or long runs of DNA segments shared between individuals, as being regions of ibd (Gazal *et al*. 2014, Browning and Browning 2012, Ringbauer *et al*. 2024) as we also discuss below. First, though, we review allele-frequency based methods.

For population *i*, the sample frequencies 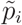 for the designated alleles are one half the average of the sampled individual allele dosages 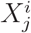. We refer to the variation of 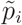 about the population frequency *p*_*i*_ as statistical variation (Weir 1996). Each *p*_*i*_ represents a single realization of an evolutionary process and the average over all realizations, and all populations, is the allele probability *π*. We refer to the variation of the *p*_*i*_ about the probabilities *π* as evolutionary or genetic variation (Weir 1996). All instances of a variant in a study are assumed to have the same probability *π* and we consider this to be a nuisance parameter. Although *π* can be estimated by any sample allele frequency 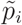, nonlinear functions of 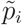 have expected values affected by both statistical and genetic sampling. The issue is that unrecognized ibd in a sample reduces the number of independent copies of an allele below what might be expected from the sample size. From Equation 1 and the expressions in Box 1, the expected value of 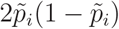, for example, is (Weir 1996)

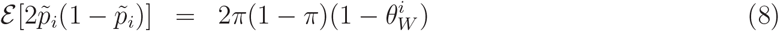

From the definition of 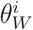 in Box 1, this expectation depends on the number *n*_*i*_ of individuals in the sample. Since 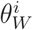 is not negative, the right hand side of Equation 8 is always less than 2*π*(1 − *π*). As 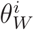 is unknown, there is no unbiased estimator of 2*π*(1 − *π*) although some authors have implied that 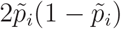 is such an estimator if there is no inbreeding (Manichaikul *at al*. 2010), or if the population is in Hardy-Weinberg equilibrium (Patterson *at al*. 2006) or if the sample size is large (Milligan 2003). In all these cases the estimator fails to be unbiased for 2*π*(1−*π*) because the descent measure 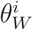 is unlikely to be zero. The proposal by other authors (DeGiorgio and Rosenberg 2009) to estimate 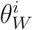 from pedigree data in order to provide an unbiased estimator does not account for pedigree-based ibd probabilities being relative to the pedigree founders and being expected values that might not represent the actual ibd status. The inclusion of the sample size in measure 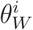 illustrates the dependence on study dimensions of properties of estimators that make explicit use of sample allele frequencies. A good discussion of the issues raised by using sample allele frequencies has been provided by Wang (2014, 2017, 2022).

Wright’s publications, including Wright (1943, 1965), involved variances of allele frequencies over populations and often made use of heterozygosities. His work was the basis for an estimator of the within-population inbreeding coefficient given by Li and Horvitz (1953). Those authors implied they were estimating the population-specific inbreeding coefficient 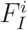 value, but gave an estimator for the population-specific 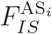. By manipulating Equation 1, and replacing 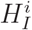 by the observed proportion 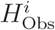 of heterozygotes in population *i*, their estimate was derived to be

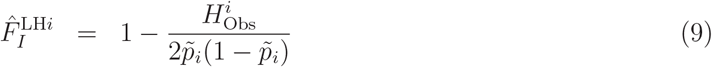

This estimator has been presented by many authors since 1953, e.g. Reich *et al*. (2009). Although 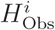 is unbiased for the probability 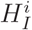 an individual in population *i* is heterozygous, Equations 1 and 8 show that, if the expected value of the ratio can be approximated by the ratio of expectations and if the sample size is large, the Li and Horvitz estimator is for 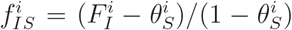, not 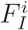. The ratio form of the estimator allows the unknown parameter 2*π*(1 − *π*) to cancel out of its expectation, but there is still the inherent non-estimability of the descent coefficient 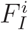 in the absence of information about a reference population in which there is known or zero correlation or identity by descent. Moreover, the necessity of assuming large sample sizes makes this estimator of limited value for many studies of natural populations and this example informs our recommendation 4 not to use estimators that involve sample allele frequencies. We return to the estimators of Nei, Cockerham, VanRaden, Yang and others later in this discussion.

#### Allele-Sharing Methods

A more recent approach (Rousset 2007; Speed and Balding 2015; WG17, Goudet *et al*. 2018; Ochoa and Storey 2021 (OS21 hereafter); Zhang *et al*. 2022; HO23; Goudet and Weir 2023 (GW23 hereafter) does not use sample allele frequencies and is designed to follow the reformulation of Wright’s *F*-statistics given in the Introduction. The unobservable descent status for a pair of alleles is replaced by observable allele-pair identity in state. As in Rousset (2007) we quantify allele-sharing as the proportion of pairs of distinct alleles that have the same state. There is only one pair within an individual and the proportion is 1 for homozygotes and 0 for heterozygotes. There are four pairs for two individuals and the proportions are 1 when they are both homozygous for the same allele, 0 when they are both homozygous but for different alleles, and 0.5 for all pairs involving heterozygotes.

For individuals, the allele-sharing statistics *Ã* are expressed in terms of allele dosages in Box 2, along with various averages. For pairs of individuals, we can also use individual-specific sample allele frequencies 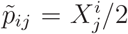 for individual *j* in population *i*. Those expressions for 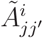 have the same form as 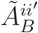 for pairs of populations *i, i*′ and both statistics are the same when population samples reduce to a single individual, *n*_*i*_ = 1.

Box 2

Briefly, there is an allele-sharing statistic *Ã* for every *θ* parameter in Box 1 and estimation rests on a consequence of Equation 1. For any allele pair(s)

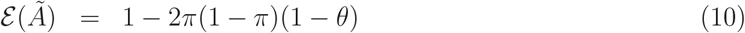

Estimators for the descent measure for allele pairs *Y* relative to allele pairs in set *R* are

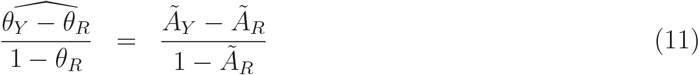

These estimators describe the dependency between allele pair(s) *Y* relative to the average dependency over all pairs in *R*. As *R* is the same for all members of *Y*, the estimator expectations have the same ranks as do the *θ*’s (Recommendation 2).

The estimators in Equation 11 can be applied to all the measures in the left hand sides of Equations 2-5 by replacing the *θ*’s on the right hand sides by *Ã*’s. Our use of a tilde on *Ã* emphasizes that these are observable statistics, and we write *A* for their parametric expected values. A complete set of allele-sharing estimators is shown in Box 3 for inbreeding, kinship and population structure. Each estimator 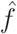 or 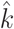 has an expected value of *f* or *k* given by replacing the *Ã*’s by *A*’s. The *A*’s, in turn, are functions of the ibd coefficients *F* or *θ* for the alleles identified in the *A* suffices. The extension of allele-sharing estimators from individuals to populations is seen to be seamless. Note that both individual-specific inbreeding and individual-pair-specific kinship are estimated relative to average kinship in the reference set (Recommendation 5): either the individuals’ own population or the whole study.

Box 3

The population-structure estimators 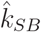, or 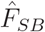, in Box 3 are the allele-sharing versions of Wright’s *F*_*ST*_. The population-specific values 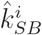 may help in detecting indications of natural selection (Weir *et al*. 2005). It has been common to present pairwise estimators of population structure (e.g. Hudson *et al*. 1992, Cumer *et al*. 2022) and these are shown in Box 3 as 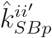 which is just the value of 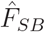 for a study with only two populations.

Multiple variants are accommodated by summing the allele-sharing statistics over variants in the numerators and denominators in Equation 11 separately before computing their ratio. Numerators and denominators both satisfy the conditions of OS21 for the ratios to have expected values that converge almost surely to (*θ*_*Y*_ − *θ*_*R*_)/(1 − *θ*_*R*_) as the number of variants increases, regardless of the number of individuals or populations (Appendix A).

The estimators 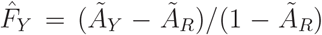 and their expectations may be negative, but the reference set of alleles *R* can always be changed to make any estimate, or average estimate, positive or zero if that is desired. Changing the reference set from *R* to *R*′ changes the estimator to 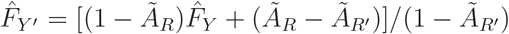. The expectations of 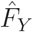 and 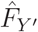 both have the same ranks as those of the target *θ*_*Y*_ ‘s. If *Y*_min_ is the value of *Y* that minimizes *Ã*_*Y*′_, setting 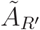 to 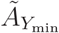 gives 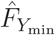 a value of zero as used by OS21 and by HO23. For no value of *R*, however, are ibd probabilities themselves being estimated as that is not possible when allele probabilities *π* are not known.

We show in Appendix B that reducing genotypic data to allelic data provides allelic association estimates that are affected by departures from Hardy-Weinberg equilibrium. Genotypic data should be used if they are available (Recommendation 3).

##### Popkin Estimators

The estimators of OS21, HO23, implemented in the Popkin package (Ochoa and Storey 2023), are broadly similar to those discussed here and their results and conclusions are also similar. There are aspects of their work, however, that can be noted.

Contrary to what is written in OS21 and HO23, the allele-sharing estimators in WG17 are unbiased, as we showed most recently in GW23. OS21 were the first to show that ratio estimators of allele-sharing statistics using large numbers of genetic variants are unbiased *for the compound parameters they were designed to estimate*. The larger point, that ibd probabilities are estimated in a “relative to” sense, has been central to descent measure characterization since the early work by Sewall Wright. More explicitly, Cockerham (1973, page 682) said

> Thus in practice the model is reduced to accommodate the parameters estimable from the information or data available, and the estimable correlations are always relative to the correlation of genes farthest apart (generally least related and least correlated) in the informational system. … points out the result of enforcing the constraint that *θ*_*B*_ = 0 [our notation] when it actually is not zero.
>
> The same point was made by Chesser (1991), ourselves (Weir and Hill 2002) and Bhatia *et al*. (2013).

OS21 and HO23 make their estimates relative to the minimum descent measure value in a reference set of alleles whereas our estimates were relative to an average value. Averages are likely to be less variable than minima, and GW23 showed (Figure S4) that confidence intervals for OS21 estimates were wider than for WG17 and GW23 estimates. There is a simple linear relationship between the estimates in OS21 and HO23 and those in WC84, WG17 and GW23. Identity by descent parameters *θ* are not being estimated in these publications.

#### Nei Estimators

Nei (1977) defined three study heterozygosities: the proportion *H*_0_ of heterozygotes for all individuals and *H*_*S*_, *H*_*T*_ for the Hardy-Weinberg-equilibrium expected proportions of heterozygotes within populations and in the total of all populations, respectively. These are expressed in terms of population allele frequencies as shown in Box S1 and were used to define *F*-statistics. His *F*_*ST*_ was also given in Nei (1973). Nei was concerned with characterizing extant populations without considering genetic sampling, so population allele frequencies were fixed. Population allele frequencies are not known, so Nei’s analyses made use of sample allele frequencies within populations and across all populations. As with Wright’s original formulation involving random pairs of alleles, basing analyses on sample allele frequencies leads to estimators with expected values depending on the numbers of sampled populations and the numbers of individuals from each population.

In particular, the estimator 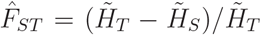 given in Nei (1973, 1977) for large sample sizes has an expected value, following Box S1, of 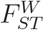 shown in Equation 4. Only for large *r* is this the value of 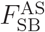 in Equations 3.

In subsequent publications, Nei and Chesser (1983) and Nei (1986), Nei modified his estimators to reduce the effects of study dimensions. We show in Box S1 that his treatment in these later publications essentially replaced his *H*_*S*_ (our *H*_*W*_ = 1 − *A*_*W*_) by our *H*_*S*_ = 1 − *A*_*S*_, and his *H*_*T*_ by our *H*_*B*_ = 1 − *A*_*B*_. The original functions of sample allele frequencies were replaced by allele-sharing statistics for distinct individuals or distinct populations. The estimators of Nei (1986) are just the allele sharing estimators of the previous section and we recommend that analyses be conducted with allele-sharing statistics (Recommendation 2).

Bhatia *et al*. (2013) also ignored genetic sampling, with population allele frequencies being fixed, but they were critical of Nei’s approach. However, their definition and estimator of *F*_*ST*_ (their equations (7) and (s13)) are not the definition and estimator given by Nei (1986). It may prevent confusion to use Nei (1986) when referring to “Nei’s Estimators” and these estimators are also given in Nei’s textbook (Nei 1987) where he mentions the advantage of not working with estimators that depend on the number of populations. Initially Nei (1973, 1977) defined *F*_*ST*_, in our notation, as (*A*_*W*_ − *A*_*T*_)/(1 − *A*_*T*_) and then in Nei (1986) changed to our (*A*_*S*_ − *A*_*B*_)/(1 − *A*_*B*_). These expressions do not support the claim (Bhatia *et al*. 2013; Gusev 2024) that Nei’s two estimators differ by a factor of 2 when there are two populations.

#### Cockerham Estimators

Wright’s original formulation of *F*-statistics was in terms of correlations of pairs of alleles, and Cockerham (1973) also used correlations. Cockerham expressed allelic indicators as linear models with a mean and effects for populations, for individuals within populations and for alleles within individuals. The components of the total variance *π*(1 − *π*) for an indicator were written as *a, b, c* respectively for these three effects. The parametric forms of the *F*-statistics can be written as intra-class correlations with estimators using variance-component estimates: 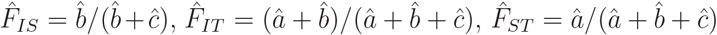.

WC84 used an analysis of variance (anova) to set out the estimation of variance components and correlations. They used sample-size weights to combine statistics over populations and an algebraic comparison with the model of WG17 is cumbersome, but when sample sizes are equal for all populations there is algebraic identity between the *F*-statistic estimators of the two approaches. (Boxes S2 and S3).

WC84 assumed there was no ibd between sampled populations and that all populations had the same values for descent measures. Under those assumptions, their mean squares are identical to allele-sharing expressions (Boxes S2, S3) for all sample sizes and lead to the descent measure estimates predicted by Cockerham (1973). We would prefer not to make those earlier assumptions so we recommend the allele-sharing approach shown in WG17 without weighting over populations by sample sizes (Recommendations 2, 4), rather than the approach in WC84. Rousset (2007) also pointed out the equivalence of the allele-sharing and Cockerham estimators for equal sample sizes.

For allelic data the analysis of variance has only two sources of variation: between and within populations. Only *F*_*SB*_ can be estimated in the absence of genotypic information and the estimator given in WC84 has an expectation equal to the allele-sharing estimator for allelic data if sample sizes are all equal or if ibd probabilities are equal for all sampled populations, *and if Hardy-Weinberg equilibrium is assumed*. Otherwise, it has an expectation that depends on inbreeding and coancestry levels, sample sizes and the number of populations sampled. In general, the analysis of allelic data seems to be accomplished better by comparing allele sharing for random pairs of distinct alleles within populations (*Ã*_*D*_) to random pairs of distinct alleles between populations (*Ã*_*B*_) as shown in Appendix B. A more accurate account of population structure requires genotypic data to remove the effects of population-specific inbreeding and relatedness (Recommendation 3). There are situations of course where allelic data are not available, such as with aDNA where genotype calls are often imprecise. We suggest a way to handle this type of data in the missing data section of the Discussion.

#### Standard Kinship Estimators

Prior to the availability of extensive molecular markers for quantitative genetics, it was usual to use path counting methods (Wright 1922) to predict inbreeding and coancestry coefficients. If individuals *j, j*′ had a common ancestor *J* and there were *n* individuals in the chain *j*. ..*J*. .. *j*′ then

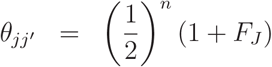

The right hand side is summed over all paths joining *j* to *j*′ through *J* and all common ancestors *J*. The coancestry of *j, j*′ is also the inbreeding coefficient of any individual for which they are parents.

Yu *et al*. (2006) were the first to replace pedigree-predicted NRM’s with those estimated from molecular markers. They pointed out that pedigree records may not be available or may be inaccurate. It is also the case that pedigree values are limited by the pedigree, and are relative to the available founders for that pedigree. They are calculated ibd probabilities and will differ from the actual ibd levels in the individuals in a study (Visscher *et al*. 2006; Hill and Weir 2011; Goudet *et al*. 2018). Marker-based estimates do not have these limitations and any valid estimation method can be used. In the quantitative genetics field, estimators using allele dosages and sample allele frequencies are commonly used (Yang *et al*. 2010, 2011; VanRaden 2008). They have been referred to as “standard” (WG17; OS21; Zhang *et al*. 2022; HO23).

The standard estimators make use of the matrix ***M*** with elements 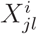, the population-*i*, individual-*j* allele dosage for variant *l*, in row *j* and column *l*. Patterson *et al*. (2006) pointed out that mean of column *l* of ***M*** is equal to 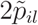, twice the sample allele frequency for variant *l* and population *i*. If 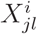 could be regarded as having a binomial distribution Bi(2, *π*_*l*_), requiring HWE in the population, it would have variance 2*π*_*l*_(1 − *π*_*l*_) with a plug-in value of 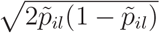. Patterson *et al*. normalized 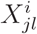 to 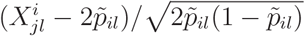 and then formed the matrix 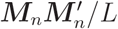 where *L* is the number of variants and ***M***_*n*_ is the matrix of normalized allele dosages. In essence, this appears to be a sample correlation matrix for individual allele dosages. Patterson *et al*. recognized the limitations of the HWE assumption, but (using our notation) Yang *et al*. (2011) stated “the genetic relationship between individuals *j* and *j*′ can be estimated by”

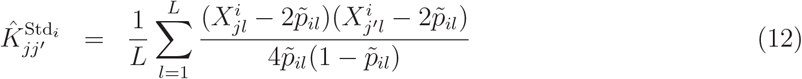

This expression applies to *j*′ being the same or different to *j*. The factor 4 in this expression is because it is the kinship matrix ***K***, rather than the GRM ***G*** being estimated.

Equation 12 employs an average of ratios over variants, even though this is well-recognized as being unstable because of potentially low sample allele frequencies in the denominator. It was shown in OS21 not to tend to its parametric value as the number of variants increases, and it is preferable to use a ratio of averages as discussed above and used in WC84, the second method of VanRaden (2008), Bhatia *et al*. (2013) and WG17. The following comments apply to the ratio of averages version.

The standard estimators are not unbiased for either 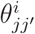 or 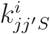 since they have large-sample expected values (WG17, Goudet *et al*. 2018; OS21; HO23) of

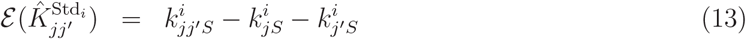

where 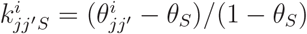 and 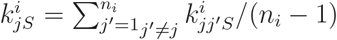. This expected value depends on the average kinship of each of the target individuals with other individuals in the sample so that the standard kinship estimates have expectations that do not have have the same rankings as the coancestry coefficients. This also applies to modified inbreeding coefficient estimators such as 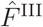 (Yang *et al*. 2011) or 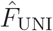 (Yengo *et al*. 2017). Neither kinship nor inbreeding, therefore, can be estimated by standard methods without considering the other (Recommendation 5).

We have shown (Box S1) that the *n*_*i*_ × *n*_*i*_ matrix 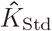 of standard estimators, for large sample sizes, is the double-centered version of the corresponding allele-sharing matrix 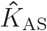 described in Box 3:

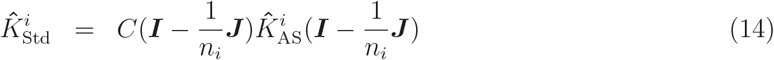

Here ***I*** is the *n*_*i*_ × *n*_*i*_ identity matrix, ***J*** is the *n*_*i*_ × *n*_*i*_ matrix of 1’s and 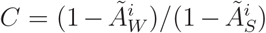. For large samples, 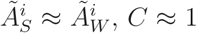, and Equation 14 leads to Equation 13. Equation 14 also holds if 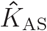 is replaced by the allele-sharing matrix *Ã*^*i*^ with elements 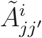 in row *j*, column *j*′ and *C* = 1.

Method 1 of VanRaden (2008) also uses the standard matrix 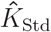 except that the sample allele frequencies “should be from the unselected base population rather than those that occur after selection or inbreeding.” This suggests those frequencies are the allele probabilities required to relate observed genotypic data to allelic correlations or ibd probabilities, as in Equation 1. If that was the case then VanRaden would be estimating ibd probabilities. However, even when information from a base population is available, as in managed plant or animal populations, the estimates will still be relative to that population. As they are predicted measures of identity by descent, they can differ from actual ibd status indicators.

#### Other Estimators

We have restricted our attention to estimation of descent measures for pairs of alleles but those may not allow distinctions among different classes of relatives: full-sibs and parent-offspring pairs, for example, have the same coancestry value of 0.25. This has led to the three-parameter set of Cotterman coefficients described by Thompson (1975). These are the probabilities two non-inbred individuals share 0, 1 or 2 pairs of ibd alleles. Fifteen coefficients are needed to describe the relatedness of inbred pairs of individuals (Jacquard 1972), although when there is no need to distinguish between maternal and parternal origins of alleles Jacquard showed a condensed set of nine coefficients.

Because real populations have only a finite number of individuals in any generation, we cannot assume a lack of inbreeding and coancestry, and recent empirical surveys (deBoer *et al*. 2021) support that assertion. Therefore, we do not review the considerable literature on the estimation of the three Cotterman coefficients (Thompson 1975; Ritland 1996) where inbreeding is not considered. We also do not review here the growing literature on the estimation of Jacquard’s coefficients (Milligan 2003; Csűrös 2014; Wang 2022; Graffelman *et al*. 2025) since those papers use sample allele frequencies and cannot, therefore, be estimating ibd probabilities. Wang (2017) reviewed many of the estimators in common use at that time to show the effects of using sample frequencies from the same populations for which relatedness estimators were sought. Thompson (1975) discussed the two issues of inbreeding and unknown allele probabilities and a full treatment is beyond the scope of this review.

##### KING

We do note the work of Manichaikul *et al*. (2010) on relatedness estimation because they considered not using sample allele frequencies. Their KING-robust estimator uses individual heterozygosities for two target individuals *j, j*′ instead of sample allele frequencies 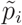 from all individuals in a sample from population *i*. In our notation their estimator is

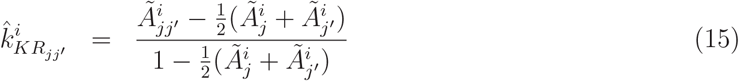

where the allele-sharing statistics can be expressed in terms of individual-specific sample allele frequencies (Box 2). Moving from whole-sample frequencies 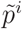 to 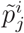 does not avoid the confounding of the expected values of their estimator by other ibd probabilities (inbreeding coefficients). The expected value of 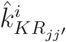 follows from Equation 10 and was given in Appendix 2 of Conomos *et al*. (2016):

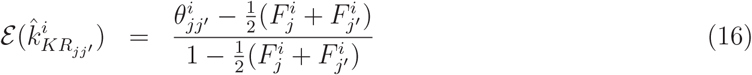

Although setting 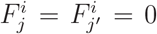 suggests that 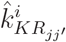 is unbiased for 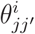, it is actually estimating coancestry relative to the average inbreeding for *j, j*′.

Manichaikul *et al*. (2010) see it as an advantage that their robust estimator “uses only information from the two individuals under comparison” but this seems to us to be a disadvantage. To paraphrase Thompson’s earlier quotation, there is no absolute measure of coancestry: the degree of relationship of any pair of alleles from two individuals can be assessed only by comparison with other pairs of alleles, possibly from within individuals, but preferably from other pairs of individuals. The complex relationship between inbreeding and kinship means that the estimator in Equation 15 is unlikely to rank pairs of individuals according to their coancestry values, and we confirm that empirically in the Results.

##### Hudson

For allelic data, Hudson *et al*. (1992) estimated *F*_*ST*_ with their statistic 1 *−H*_*w*_*/H*_*b*_ and their verbal definitions indicate this was the allele-sharing estimator for allelic data. Although those authors did not give expressions for calculating *H*_*b*_, *H*_*w*_ from data, Equations (s17), (s18) of Bhatia *et al*. (2013) and Equations (5), (6) of Mary-Huard and Balding (2023), confirm that the sample values are 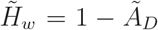 and 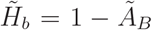. Those authors consider only two populations but the expressions hold for other numbers. Because all these authors were using allelic rather than genotypic data they were estimating 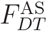 rather than 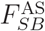. As they state, their estimates are for 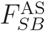 if inbreeding and coancestry are equal in every population (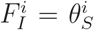, Appendix B). Reich *et al*. (2009) and Patterson *et al*. (2012) also presented the Hudson estimator for allelic data. Reich *et al*. added genotypic data to give the allele-sharing estimator 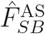 that is “robust to inbreeding” as shown in Box S4.

##### *f*-statistics

Reich *et al*. (2009) introduced a trio of “*f*-statistics” *f*_2_, *f*_3_, *f*_4_, designed to characterize population admixture by estimating population-pair coancestries 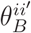 and they gave further details in Patterson *et al*. (2012). They framed their development in terms of actual allele proportions in populations, as did Nei (1973), and like Nei (Nei and Chesser 1983; Nei 1986) their estimators can be expressed in terms of allele-sharing, as we show in Box S4. For all sample sizes their estimators “in the presence of inbreeding” (Supplementary Information of Reich *et al*. 2009) do not require Hardy-Weinberg equilibrium in each population (they do not assume 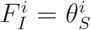) and have expectations

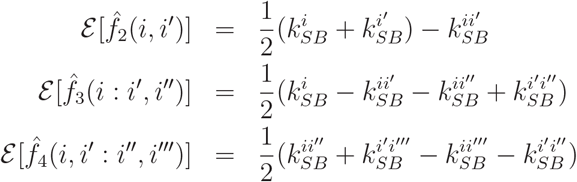

where 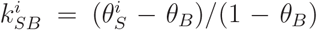 and 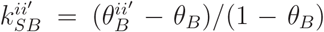 (Box S4). For independent populations, 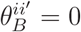 for all *i, i*′ and the expression for *f*_3_(*i* : *i*′, *i*′′) reduces to the population-specific value 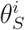 as noted by Shriver *et al*. (1997). It is not immediately clear from these expectations that a negative value of *f*_3_(*i* : *i*′, *i*′′) “provides unambiguous evidence of population mixture in the history of population *i* [our notation]” (Appendix B of Patterson *et al*. 2012).

Using a common denominator for *f*-statistics over sets of two, three or four populations allows the values to be compared although using 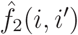 as a (drift) distance measure for populations *i, i*′ generally employs a different denominator for each pair of populations (Reynolds *et al*. 1983), as shown by estimators 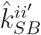 in Box 3. We note 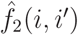 is not of the form of equation 11 and is not an unbiased estimator of 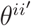.

To prevent confusion with “*F*-statistics” referring to the trio *F*_*IT*_, *F*_*IS*_, *F*_*ST*_ (Wright 1951) we suggest that the *f*-statistics of Reich *et al*. (2009) and Atag *et al*. (2024) not be termed *F*-statistics (Peter 2016).

##### Shared Segments of Identity by Descent

The methods we have discussed lead to statements about identity by descent based on observed identity in state. Another class of estimators, termed “Shared ibd segments” or “Runs of homozygosity” focus on genomic regions that are thought to be ibd. As discussed by Gibson *et al*. (2006) an individual homozygous for many variants in an uninterrupted sequence is likely to be autozygous in that region, meaning that the same chromosomal segment has been passed to the individual from each parent who inherited it from an ancestor they have in common. Under that assumption, the segments may be referred to as being identical by descent. For pairs of individuals, the corresponding procedure is to search for shared long haplotypes: this requires phased data, although single-SNP genotype methods were described by Henn *et al*. (2012). Those authors required individuals to share at least one allele for each SNP in a region. Recombination breaks up long segments over time, so long ibd regions can indicate identity by descent to an extent that depends on the number of generations back to parental common ancestors. The proportion of the genome inferred to be in regions of autozygosity is taken to be an estimator for the individual inbreeding coefficient or individual-pair coancestry. Gibson *et al*. (2006) point out that long ibd segments runs can also arise from other mechanisms, including mutation, uniparental disomy, low recombination and linkage disequilibrium, deletions and selection. Long ibd segments result in shared haplotypes across populations, as reported by Conrad *et al*. (2006) and Leslie *et al*. (2015).

Determining how long a region should be before it can be designated as ibd is subjective, and several publications give strategies for identifying regions where ibd is assumed (Gazal *et al*. 2015; Joshi *et al*. 2015). These observational strategies include adjusting the size of windows in which SNPs are characterized, overlap between windows, the numbers of heterozygotes or no shared alleles allowed per window, the total length of ibd regions and so on. These software parameters affect the size of the estimates (Meyermans *et al*. 2020). Lavanchy *et al*. (2023) discussed the effects of using reduced genomic representation, such as provided by SNP arrays, in declaring a region to be ibd. If variants within such a region are not in the reduced data, they may be polymorphic and the length of inferred ibd regions will be overestimated. Alternatively, model-based methods (Gazal *et al*. 2015; Narasimhan *et al*. 2016; Druet and Gautier 2017, 2022), using hidden Markov models where ibd is the hidden status of an observed homozygote, necessarily have assumptions such as knowing allele probabilities and there being HWE in the sampled population. The models of Druet and Gautier allowed for different classes of homozygous runs, corresponding to different times when the parents of a target individual had a common ancestor.

Estimation of apparent identity by descent within or between individuals seems to contradict Thompson’s observation that ibd is not absolute. This contradiction is resolved by noting these methods report results relative to the parameters set to define when a region is long enough to be called ibd and do not involve a reference set of individuals. Shorter threshold lengths allow for more distant common ancestors to the target individuals and the lengths might be adjusted to distinguish between recent (familial) ibd and distant (evolutionary) ibd. Allele-sharing estimators do not make this distinction. Shared ibd segment methods do not pay attention to our central point that ibd, and hence degree of relatedness, is context dependent. As argued eloquently in Chapter 8 of Walsh *et al*. (2025), estimation should specify the search parameter values used to identify ibd regions and present the fraction of the genome in those regions rather than making a designation of specific classes of relationship such as specific cousinships.

There is little doubt that ibd-region methods have had a significant impact on many genetic analyses, including the recent field of forensic genetic genealogy (Dowdeswell 2022). Descendants in common among people identified as being related to the source of an evidential sample, even if the exact class of distant relationship may not be correct, can point to one or very few individuals likely to be that source.

##### Ancestral Recombination Graph (ARG)

The methods reviewed so far have been descriptive: they characterize the sampled populations without needing to invoke details of evolutionary processes such as mutation and recombination. There has recently been substantial work done with ancestral recombination graphs (ARGs) where those processes are considered explicitly (e.g. Tsambos 2022; Harris 2023; Zhang *et al*. 2023; Wong *et al*. 2024; Lehmann *et al*. 2025). ARGs describe the network of inheritance relations among a set of individuals that result from mutation and recombination (Lehmann *et al*. 2025) and are used to describe relatedness as the total area of the branches in the ARG ancestral to both individuals (Fan *et al*. 2022). This is a multi-locus analysis and allows relatedness to be quantified at different times in the past. The computational requirements are formidable but efficient methods are appearing (Zhang *et al*. 2023; Wong *et al*. 2024).

ARG methods are based on gene genealogies for the haplotypes carried by sets of individuals and the distinction between statistical and genetic sampling, with the associated variances of sample allele frequencies, is not generally made. The “trait-centric” notion of genetic relatedness (Lehmann *et al*. 2025) appears to be equivalent to the standard kinship estimation procedure, and their “trait-centric perspective on branch relatedness” appears to have the same issue of individual-pair relatedness being confounded by the relatedness among all pairs in the sample we noted for standard estimates. A recent and comprehensive review of ARG methods was given by Nielsen *et al*. (2025).

## Results

We have previously (WG17; Goudet *et al*. 2018; GW23; Lavanchy *et al*. 2024), reported descent measure estimates for 1000 Genomes data (The 1000 Genomes 2020), using allele-sharing and other estimators. Those estimates, and those we now show, were produced with our *hierfstat* software (Goudet 2005).

For new results we concentrate on context: the need to consider both inbreeding and relatedness in characterizing either and the need to specify reference sets of alleles. We have used 61,599,150 bi-allelic SNPs over the 22 autosomes of the high coverage version of the 1000 genomes project (Bryska-Bishop *et al*. 2022). For individuals *j, j*′ in continental area *i*, we expect the allele-sharing kinship estimators 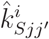 to have the same rankings as the ibd probabilities 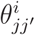, but the KING-robust estimators to be affected by the inbreeding coefficients of *j* and *j*^*×*^. This is confirmed in Figure 1. Although there is a general positive correlation between the two sets of estimates in each continental area allowing the distinguishing of close relatives from non-kin, there is also considerable variation in their ranks in some areas, as expected by our previous finding (Zhang *et al*. 2022) of a complex relationship between inbreeding and kinship values. The distinct group of negative KING-robust estimates from the AFR continental area corresponds to individual NA20314 from the ASW population who has an allele-sharing inbreeding estimate 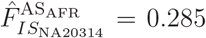. The distinct groups of negative 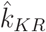 estimates for the SAS continental area reflect inbreeding levels of one or both individuals in each pair. For instance, individual HG04070 from the ITU population has an allele sharing inbreeding coefficient of 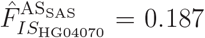. Its KING-robust kinship estimates with other SAS individuals range from 0.09 to 0.21. The distinct smaller group of negative allele-sharing kinship estimates for the AMR area are for pairs with one individual with a negative inbreeding coefficient, suggesting admixture or a different ancestral group for these individuals. The bulk of negative KING-robust estimates for AMR are due to 17 inbred individuals 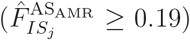 from the PEL population (the first five of which are HG02272, HG01926, HG01920, HG01961, HG01938), whose KING-robust kinships with other AMR individuals are all negative.

**Figure 1:**
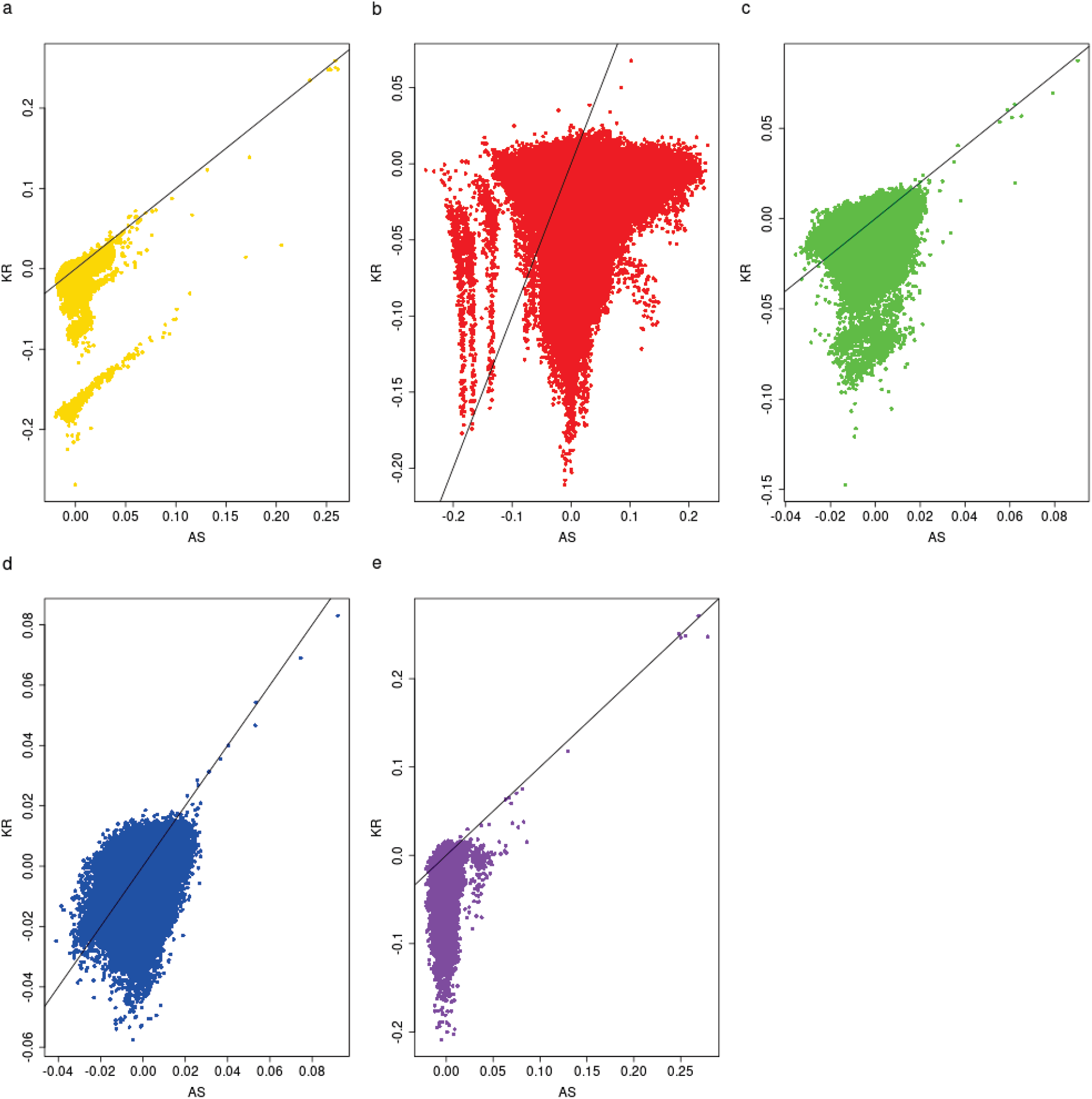
KING-robust estimates of pairwise kinship estimates 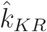 on the Y-axis versus allele-sharing estimates 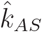 on the X-axis for 1000 Genome data from five continental-ancestry groups. a: Africa (AFR, Gold); b: Americas (AMR, red); c: East Asia (EAS, green); d: Europe (EUR, blue); e: South Asia (SAS, purple).

We repeat the analysis in Figure 1 after restricting attention to individuals whose allele-sharing estimate 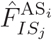 is less than 0.04. The concordance in ranks between allele-sharing and KING-robust estimates, shown in Figure 2, is now much higher.

**Figure 2:**
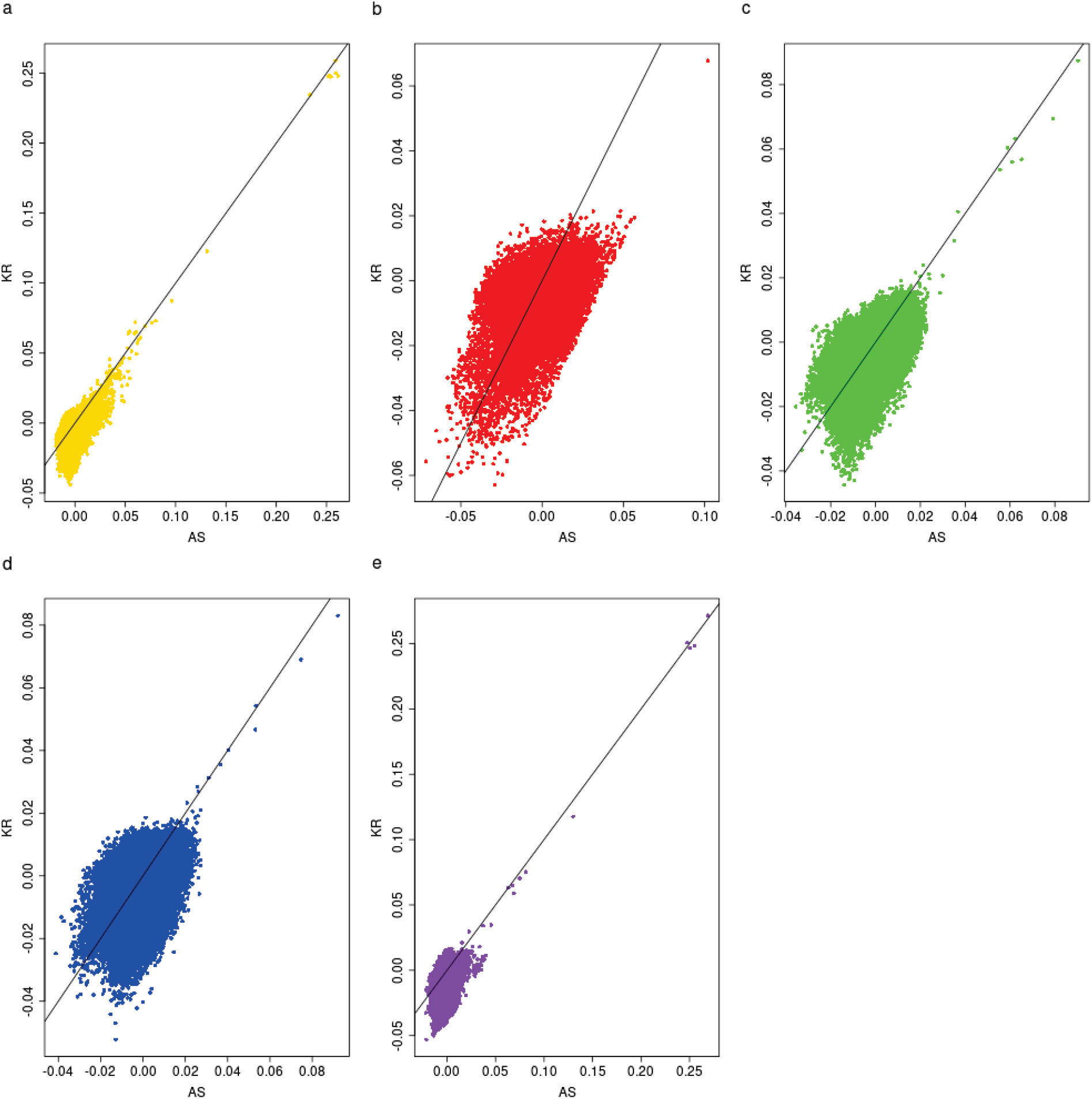
KING-robust estimates of pairwise kinship estimates 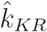 on the Y-axis versus allele-sharing estimates 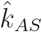 on the X-axis for 1000 Genome data from five continental-ancestry groups, using only individuals with an allele-sharing inbreeding estimate 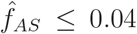. a: Africa (AFR, Gold); b: Americas (AMR, red); c: East Asia (EAS, green); d: Europe (EUR, blue); e: South Asia (SAS, purple).

Figure 3 shows a heatmap of kinship estimates from the 1000 Genomes (Bryska-Bishop *et al*., 2022). The upper half matrix shows the allele-sharing kinship 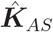 while the lower half matrix shows the standard kinship 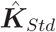. These kinships were estimated using all the pairs from the whole world as a reference. The 2504 samples are ordered by continents (AFR, AMR, EAS, EUR and SAS), and by populations within continents. African pairs all have negative allele-sharing kinships, but large positive standard kinships (larger than most non-African pairs). Since African genomes are the most diverse, we do expect they differ more one from the other than a random pair of individuals from the world and hence have a negative kinship relative to a random pair from the world. Pairs from East Asia show the largest allele-sharing kinship on average, and this matches well with their lower genomic diversity. Pairs of individuals, one from Europe and one from Asia show positive allele-sharing kinship but negative standard kinship, despite European and East Asian sharing the same set of ancestors from when humans left Africa and colonized the world.

**Figure 3:**
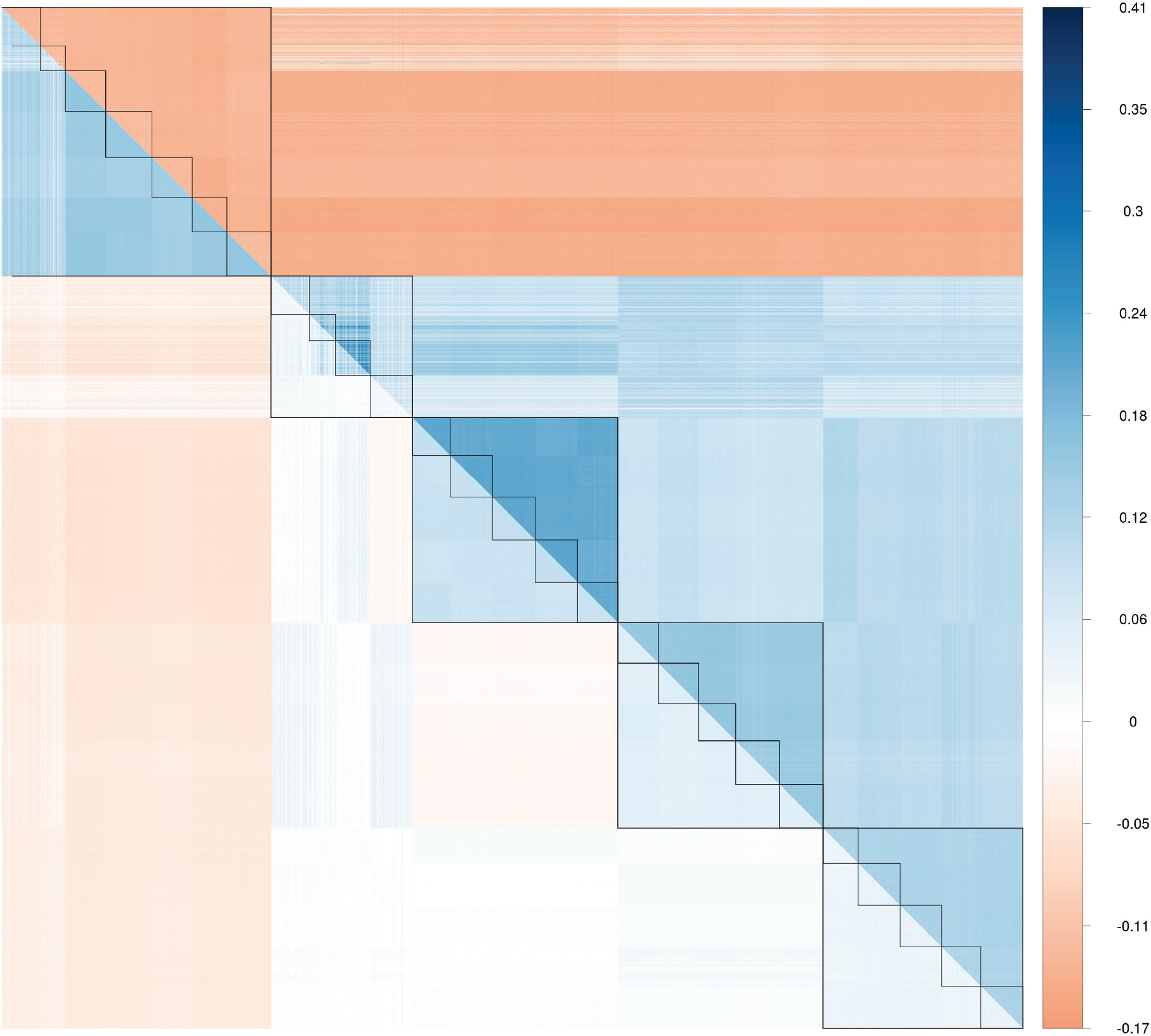
Heatmap of allele-sharing kinships 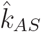 (upper half matrix) and standard kinships 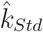 (lower half matrix) for the 2504 samples from the 1000 genomes. Inner lines delineate populations, outer lines continents. Continents and populations are in the following order (left to right and top to bottom): Africa: ACB; ASW; ESN; GWD; LWK; MSL; YRI. Americas: CLM; MXL; PEL; PUR. East Asia: CDX; CHB; CHS; JPT; KHV. Europe: CEU; FIN; GBR; IBS; TSI. South Asia: BEB; GIH; ITU; PJL; STU.

We have also looked at the effect of reference population on individual-pair allele-sharing kinship estimates, using the same data described in Bryska-Bishop *et al*. (2022). We used either the continental area or the whole world as reference for each pair of individuals within each of the 26 populations. There was perfect concordance across reference sets for the allele-sharing estimate ranks (see the left panel of Figure S1 for an illustration). For the standard estimates there was concordance for the European and Asian populations, as shown in Figure 4, but for the admixed American populations there was much less concordance between the ranks from the two reference sets and essentially no concordance for the admixed African populations. This reflects the prediction in Equation 13 that standard kinship estimates are affected by the coancestry of each pair pair of individuals as well as the average coancestry of each of the two individuals with all other individuals in the reference set: different reference sets will provide different rankings for kinship estimates.

**Figure 4:**
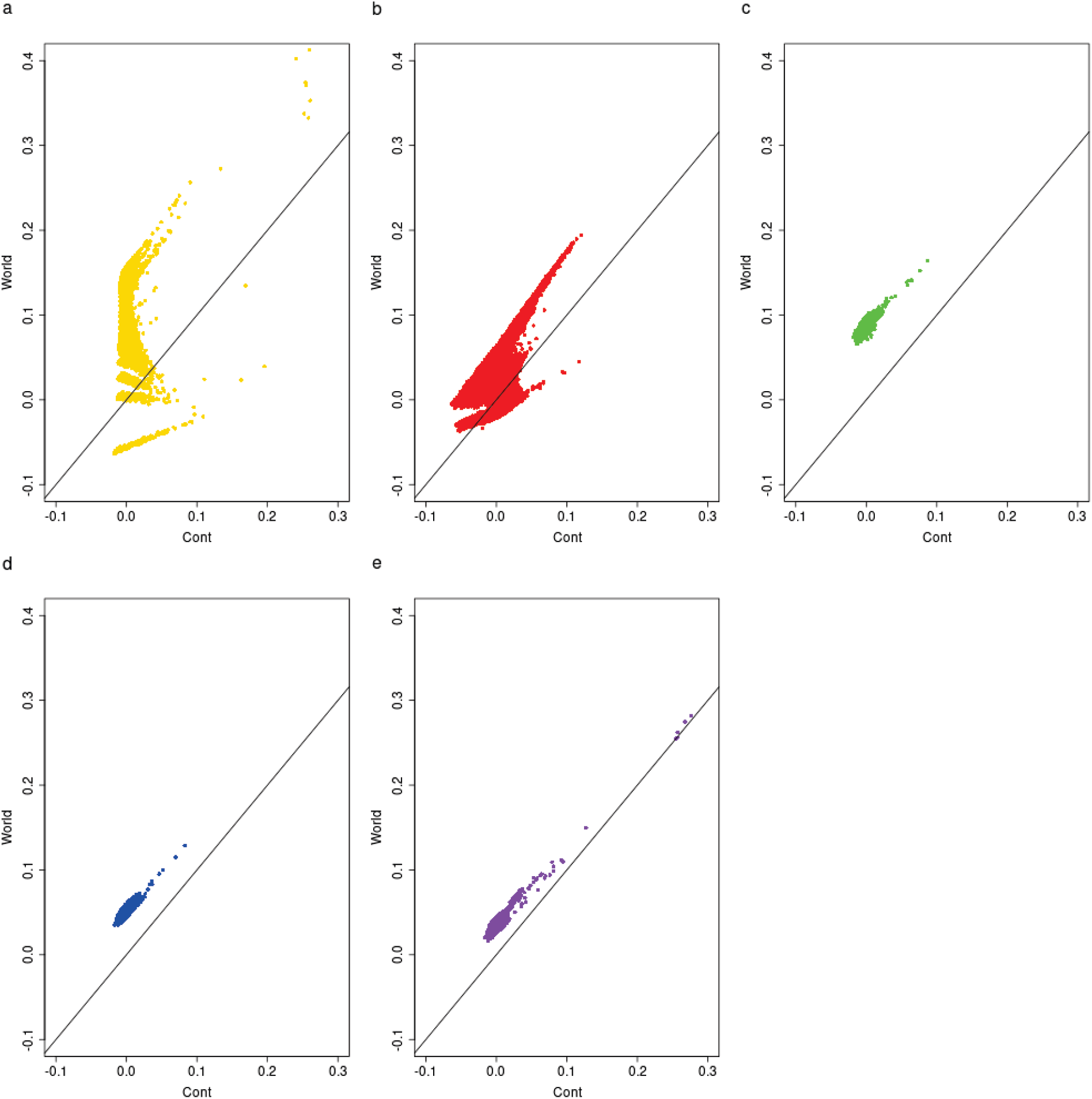
Standard individual-pair kinship estimates 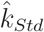, Equation 14, with either World (Y axis) or Continent (X axis) as reference population. a: Africa (AFR, Gold); b: Americas (AMR, red); c: East Asia (EAS, green); d: Europe (EUR, blue); e: South Asia (SAS, purple).

Figure S1 looks at the African samples in more detail and shows that, as expected, the allele-sharing kinship maintains the ranking of kinships when the reference is changed from the African continent to the world, whereas the standard kinship does not and reverses sometimes the rankings, as exemplified by pairs of individuals belonging to the ASW (African Ancestry in South West USA, in red) population.

## Discussion

We have looked in detail at several widely-used estimators of descent measures, with a focus on the reference set of alleles in formulating allele-sharing (AS) estimators for ibd probabilities relative to values for that reference set. The principal difference between AS and the classical Wright population=level estimators is that both target and reference sets of alleles are from distinct individuals or populations, rather than being drawn randomly. The earlier WC84 estimators are unbiased under their stated assumption that data were sampled from populations evolving independently under identical evolutionary scenarios. Cockerham (1973) pointed out that such estimators 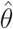 were actually estimating (*θ* − *θ*_*B*_)/(1 − *θ*_*B*_) where *θ*_*B*_ is the descent measure for alleles from different populations. Under the stated conditions, or when samples sizes are the same for all populations, the WC84 estimators are identical to the AS estimators (WG17, GW23) and we suggest that allele-sharing estimators be used for greater applicability. The AS *F*-statistics are also identical to Nei’s revised estimators (Nei 1986). OS21 and HO23 make the different assumption that the least related individuals or populations have zero relatedness so their estimators 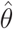 are actually estimating (*θ* − *θ*_min_)/(1 − *θ*_min_) where *θ*_min_ is the descent measure for the least related individuals or populations. Estimators based on sample allele frequencies or that assume the absence of relatedness or inbreeding may not preserve the rankings of estimates when reference sets are changed.

### Sampling Variation

Even though allele-sharing estimators are unbiased, they are subject to sampling variation. Algebraic expressions for the genetic sampling variances, for example, of these ratio estimators are cumbersome and we suggest bootstrapping over variants to give empirical values. For studies with data from large numbers of variants, observations from different variants cannot be assumed to be independent, especially for those close together in the genome. This does not affect the values and utility of AS estimators, which compare actual allele-sharing for target and reference alleles, but will affect the variance of the estimates. Block bootstrapping (Künsch 1989) divides the variants into blocks of neighboring variants and then resamples those blocks, thereby preserving within-locus dependencies. There needs to be decisions made about the size of the blocks and the degree of overlap for adjacent blocks. If descent measure estimators are to be used to identify likely causal variants for traits or targets for forces such as selection, block size needs to be large enough to reduce computational burden and reduce dependencies among blocks, and small enough to identify likely coding regions.

We provided a numerical study of variation in GW23 by simulation under a variety of population structure models and by applying block bootstrapping to analyses 1000 Genomes data. We investigated the effects of sample sizes and numbers of SNPs, and showed the expected decrease in variation as the number of SNPs increases but a much smaller effects of sample size. We found generally satisfactory performance of AS estimation with 10,000 SNPs.

### Missing Data

It is not common for a study to have every individual typed for all members of a set of SNPs and unbalanced data can have substantial implications for genetic data analyses. Graffelman *et al*. (2015) showed that spurious indications of departure from Hardy-Weinberg equilibrium can result when exact tests are applied to data with some SNPs missing from some individuals and suggested multiple-imputation to bring balance to data. Having balanced data avoids complications in determining the expected *p*-values for Q-Q plots in studies with multiple significance tests. We discussed imputation for kinship estimation in the Supplementary Information of Goudet *et al*. (2018). In our *hierfstat* package, we use all typed SNPs in an individual, or typed in a pair of individuals, and calculate allele-sharing statistics as proportions of typed SNPs rather than as proportions of all SNPs in a study. Too much missing data may introduce some bias and this issue needs further investigation.

Phased data may not be available in some studies: genotypes for each SNP are determined, but the underlying haplotype structure is not known. The AS methods we have presented here are all based on single SNPs, with simple averages being taken over SNPs. We do not assume the SNPs are independent and we do not require knowledge of phase. Our analyses of allelic data inferred from genotypic data seems appropriate for current “pseudo-haplotype” data in ancient DNA studies (e.g. Marsh *et al*. 2023). From these pseudo-haploid calls, we could calculate allele-sharing from haploid data (see next section) and estimate allele-sharing kinship. This avoids the need to estimate allelic frequencies.

### Multiple Alleles and Polyploidy

Our discussion has concentrated on loci with two alleles scored in diploid organisms but the extensions to multiallelic loci or polyploid organisms are straightforward. With only two alleles it is sufficient to focus on just one designated allele per locus as the other allele will give the same value for all estimators. To accommodate multiple alleles *u*, we just need to add subscript *u* to allelic statistics and add over values of *u*: allele dosages become 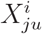, sample allele frequencies become 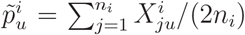, allele-sharing for individuals becomes 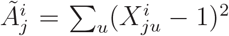 and so on. Subscripts can also be added for loci, and estimator of numerators and denominators summed over loci. Details are given in WG17.

The extension to haploids and polyploids was given by Bilton (2020) and the allele-sharing approach of WG17 with a matrix formulation suited for computation was shown in GW23. For the simple autopolyploid model reviewed by Bilton, an individual has *κ* copies of each gene and transmits a set of *κ/*2 copies chosen according to a correlated binomial distribution. The coancestry coefficient 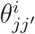 of individuals *j, j*′ in population *i* is the descent status of an allele taken randomly from each individual. There are now *κ*^2^ possible pairs of alleles and the allele-sharing statistic 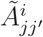 is now 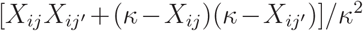 where the allele dosages can take values 0, 1,… *κ*. Although the allele-sharing statistics depend on *κ*, the allele-sharing estimators and their expectations are the same as for diploids, as shown in Box S5. There is no need to show expressions for pairs of populations as the allele-sharing statistics can then still be expressed in terms of sample allele frequencies and the estimators are still unbiased for functions of the form (*θ − θ*_*B*_)/(1 − *θ*_*B*_). Earlier treatments were given by Ronfort *et al*. (1998).

As for diploids, the descent parameter for an individual with itself depends on the inbreeding coefficient 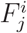 of the individual (the descent status of any two distinct alleles within the individual) and is given by 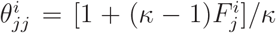. For haploids, *κ* = 1 and inbreeding is not defined and the self-coancestry is 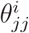 is, necessarily, equal to 1. For sample allele frequencies from three or four populations, related work was reported by Gautier *et al*. (2022).

### Relevant Parameters

Not only are compound parameters of the form (*θ*_*Y*_ − *θ*_*R*_)/(1 − *θ*_*R*_) estimable with data from extant populations, we suggest that these are actually the parameters of interest for empirical studies and we offer some examples. First, we consider the estimation of genetic distances between pairs of populations for use in the construction of phylogenetic trees. Under the simple model of random genetic drift being the only force driving the divergence of two populations from their common ancestral population, the *F*_*ST*_ value for each population decays over time at a rate that depends on population size *N*. Specifically, the ibd probability *θ*^*i*^,*i* = 1, 2 for two randomly chosen alleles from population *i*, assuming HWE, *t* generations after divergence from their common ancestral population, is

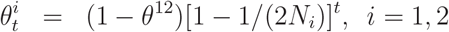

The ibd probability *θ*^12^ in the ancestral population is unknown, but it is log[(*θ*^*i*^ − *θ*^12^)/(1 − *θ*^12^)], rather than *θ*^*i*^, that changes linearly with time and can be estimated to serve as a distance. The factor *θ*^12^ was omitted by Reynolds *et al*. (1983), as they assumed between-population dependencies were zero, as in WC84. We suggested that Reynolds’ estimates for populations *i, i*^*×*^ in a set of populations be modified to 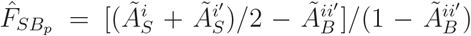 (GW23) as these estimate 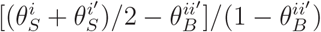 regardless of inbreeding levels.

A second example arises in forensic science, where the quantity of interest is the match probability that one person (e.g. a suspect) has a particular genetic profile given the observation of that profile in an evidential sample (e.g. from a perpetrator) under the hypothesis that the two profiles are from different people (Evett 1983). It can be expressed in terms of allele probabilities and ibd probabilities in the population of interest. It is difficult to define this population, so it seems appropriate to use average ibd probabilities such as *θ*_*W*_ that describe allelic ibd in any subpopulation of a broad population, loosely representing continental area and represented in a forensic frequency database (Ruitberg 2001). Very ofte, only allele frequencies are available from those databases. The simplest situation in this discussion is for Y-chromosome haplotype profiles *u*, where the match probability is [*θ*_*W*_ + (1 − *θ*_*W*_)*π*_*u*_]. Averaging over all profile types, the average match probability *P*_*M*_ is

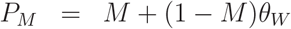

Where 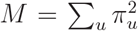. If the sample haplotype frequency 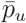 in a database is the average over many subpopulations, then the logic in Box 2 shows that sample value 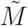 has an expected value of *M* + (1 − *M*)*θ*_*B*_, with *θ*_*B*_ referring to identity between pairs of subpopulations. An unbiased estimator of *P*_*M*_ is

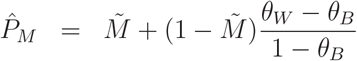

and the ratio of *θ*’s has the form of an *F*_*ST*_. Forensic scientists use recommended values, such as 0.01 (National Research Council 1996), for *F*_*ST*_ and genotypic profiles. These follow from a range of surveys and generally use the estimates from application of the methods in WC84 that we have seen are for *F*_*SB*_ rather than *θ*. It is interesting to note that on Page 103 of National Research Council (1996) the US National Research Council said

> The symbols F_ST_ (Wright 1951), G_ST_ (Nei 1973, 1977), and *θ* (Cockerham 1969, 1973; Weir 1990) have very similar meanings and for our purposes can be regarded as interchangeable.

Allele-sharing estimates of *F*_*SB*_ for forensic markers have been given by Buckleton *et al*. (2016, 2025).

Finally, in quantitative genetic studies such as the detection of inbreeding depression, the estimation of heritability and the search for causal variants that may be revealed by association mapping (GWAS). We suggest that it is the actual constellation of alleles carried by studied individuals that is of relevance, not the evolutionary mechanisms that led to those constellations. It is the deviation of target-individual actual levels of allele sharing over variants compared to the levels among reference individuals that is relevant, without regard to whether this has arisen by recent familial relatedness or by population history in the distant past. In their association studies, HO23 use their estimates of the kinship matrix rather the classical matrix of ibd probabilities and find robustness to the form of the matrix.

## Conclusions

Our conclusions are best expressed in the following set of five recommendations.

### 1: State that descent measures for pairs of alleles are relative to values in a reference set of allele pairs

At least since Sewall Wright’s work, there has been recognition that measures of inbreeding, relatedness and population structure are relative: they quantify the dependencies of specified alleles relative to dependencies in some reference population and are not absolute. Publications presenting methods of estimating these parameters, or presenting estimates from genetic data, should specify the reference population. In this paper we have generally used identity by descent as a basis for allelic dependence, but the same general principles we discuss apply also to the correlation of allelic state indicators.

### 2: Use estimators that preserve descent measure rankings over different reference sets

Along with the need to specify reference sets is the need to use estimators that preserve rankings of estimates for different target allele pairs as the reference set changes. As an extreme situation, we would not want the inbreeding estimates for two individuals to be ranked differently if the study dataset increased by adding individuals. The most dramatic example we have seen is for standard estimates showing African individuals to be the least inbred in the world with their continental area as a reference but the most inbred with all the 1000 Genomes continental areas as reference. Allele-sharing estimators do provide rank invariance of estimates. These estimates serve as descriptors of the sampled individuals and populations and do not invoke any evolutionary theories other that required for Equation 10 to hold. They are also simple to calculate and scalable to large datasets.

### 3: If genotypic data are available, avoid having to assume Hardy-Weinberg equilibrium by not reducing them to allelic data

Another of our themes has been in clarifying the role of the unit of analysis. Although descent measures usually refer to properties of sets of *alleles*, data are generally collected from individuals as *genotypes*. Genotypic data can be reduced to allelic data, but the resulting estimates are then confounded by dependencies of alleles within individuals and between pairs of individuals and may not reflect the consequences of those evolutionary forces that act at the level of individuals.

Although we have concentrated on single-SNP analyses, allele-sharing methods are best based on large numbers of SNPs and are affected by all the evolutionary forces that have led to the complete genetic profiles in the target and reference sets. Estimated inbreeding coefficients, for example, compare the actual degree of homozygosity in a target individual with the allele sharing over all pairs of individuals and this reflects the actual allelic correlations among loci in both the target and reference set. There is no need to filter datasets by minor allele frequency or linkage disequilibrium, for example. Allele sharing does not need to be independent among SNPs.

### 4: Recognize that allele frequencies do not need to be estimated

Consistent with our preference for analyses based on genotypes, we prefer not to use sample allele frequencies as these can involve alleles from the same individual, or the same population and cause study dimensions to affect resulting estimates. We show, instead, the advantage of working with genotypic measures of allele-sharing to allow a focus on only the target alleles or sets of alleles. Allele-sharing captures the descent status of alleles for specified levels within a hierarchy of individuals and populations regardless of the sample sizes at each level. Sample allele-sharing statistics are unbiased for the corresponding parametric values, and these are linear in the descent measures of interest. With data from large numbers of genetic variants, ratios of linear combinations of allele-sharing statistics are also unbiased for the parameters they are designed to estimate. They require specification of the reference set and so clarify the meaning of “relative to”.

### 5: Consider both inbreeding and kinship when estimating either one

We have stressed the need to consider both inbreeding and relatedness when coefficients for either descent status are being estimated. It is difficult to envisage a natural population with relatedness but no inbreeding for example, and our plots of kinship versus inbreeding estimates in human populations (Zhang *et al*. 2022) show that neither should be set to zero. Other authors have made the same point (e.g. Guan and Levy 2024). Some reviews of kinship estimation, however, do assume no inbreeding (Jiang *et al*. 2022).

We have shown that allele-sharing estimators satisfy our five recommendations. They are easy to calculate and interpret, and they avoid some of issues we identify with published alternatives. Even when there may be little numerical difference between allele-sharing and other estimates, we suggest using the former for a consistent approach to the analysis of genetic data.

## Data Availability Statement

We have used publicly available 1000 Genomes data, as described by Byrska-Bishop *et al*. (2022). Code to reproduce the figures is available from URL https://github.com/jgx65/GeneticsAllelicAssociation.

## Software Information

Our software package *hierfstat* (Goudet 2005) can be downloaded from URL https://cran.r-project.org/web/packages/hierfstat; https://github.com/jgx65/hierfstat.

## Acknowledgments

Very helpful comments on a previous version of this paper were received from the GENETICS Editors and Reviewers and from Sanne Aalbers and Stephen Haslett.

## Appendix A. Consistency of AS Estimators

Ochoa and Storey (2021) were the first to formalize the consistency of ratio of averages estimators for descent measures. This Appendix follows their development. The allele-sharing estimators 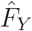 in Equation 11 are ratios of averages of allele-sharing statistics over SNPs:

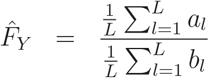

where

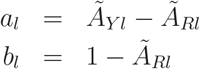

These terms have expectations

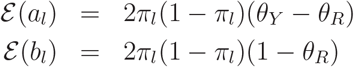

Ochoa and Storey (2021) point out that the *a*_*l*_ and *b*_*l*_ are not independent but their dependencies tend to be localized and the estimators will perform well if the effective number of independent loci is large. The *a*_*l*_ and *b*_*l*_ are not identically distributed since the *π*_*l*_ vary over loci, but each *a*_*l*_ and *b*_*l*_ is bounded above by 1 and so is its variance. It is also the case that 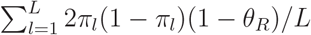 is not zero. Therefore, Kolmogorov’s criterion for the Strong Law of Large Numbers holds, based on the large effective number of SNPs, and we can conclude that 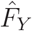 converges almost surely to (*θ*_*Y*_ − *θ*_*R*_)/(1 − *θ*_*R*_) as *L* becomes large.

## Appendix B. Allelic Data

The estimators for quantities in Equations 2-4 are for genotypic data. When genotypic data are not available there is no information in the data to allow inbreeding and coancestry to be distinguished. Distinct allele pairs within and between populations are used in 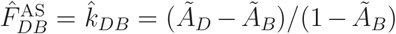 (written as *F*_*WT*_ in WG17). The expressions for *Ã*_*D*_, *Ã*_*B*_ given in Box 2 still hold for allelic data, and these lead to the expected value of 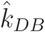:

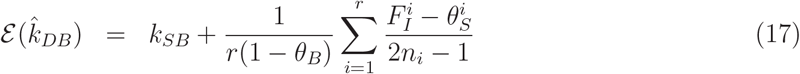

This shows that 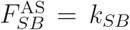 estimation from allelic data will provide the same estimate as from genotypic data either if there is HWE in every population or if every sample size is large. Otherwise the allele-based allele-sharing estimates of *k*_*SB*_ will differ from genotype-based estimates to an extent and direction that depends on departures from HWE.

### Box 1: Descent Measures^1^ for a Study with *n*_*i*_ Individuals from the *i*th of *r* Populations.

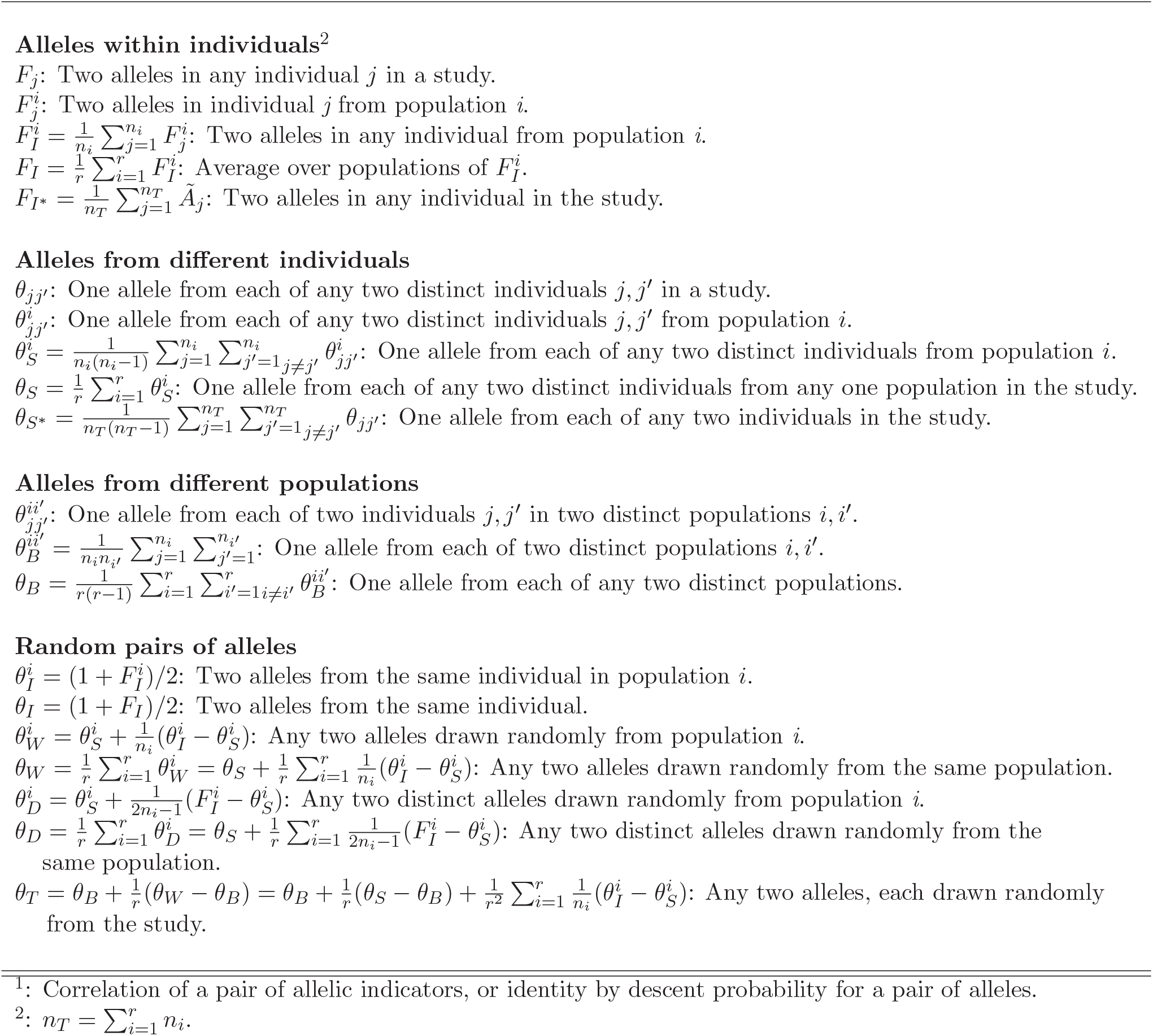

### Box 2: Allele Sharing Statistics for a Study with *n*_*i*_ Individuals from the *i*th of *r* Populations.

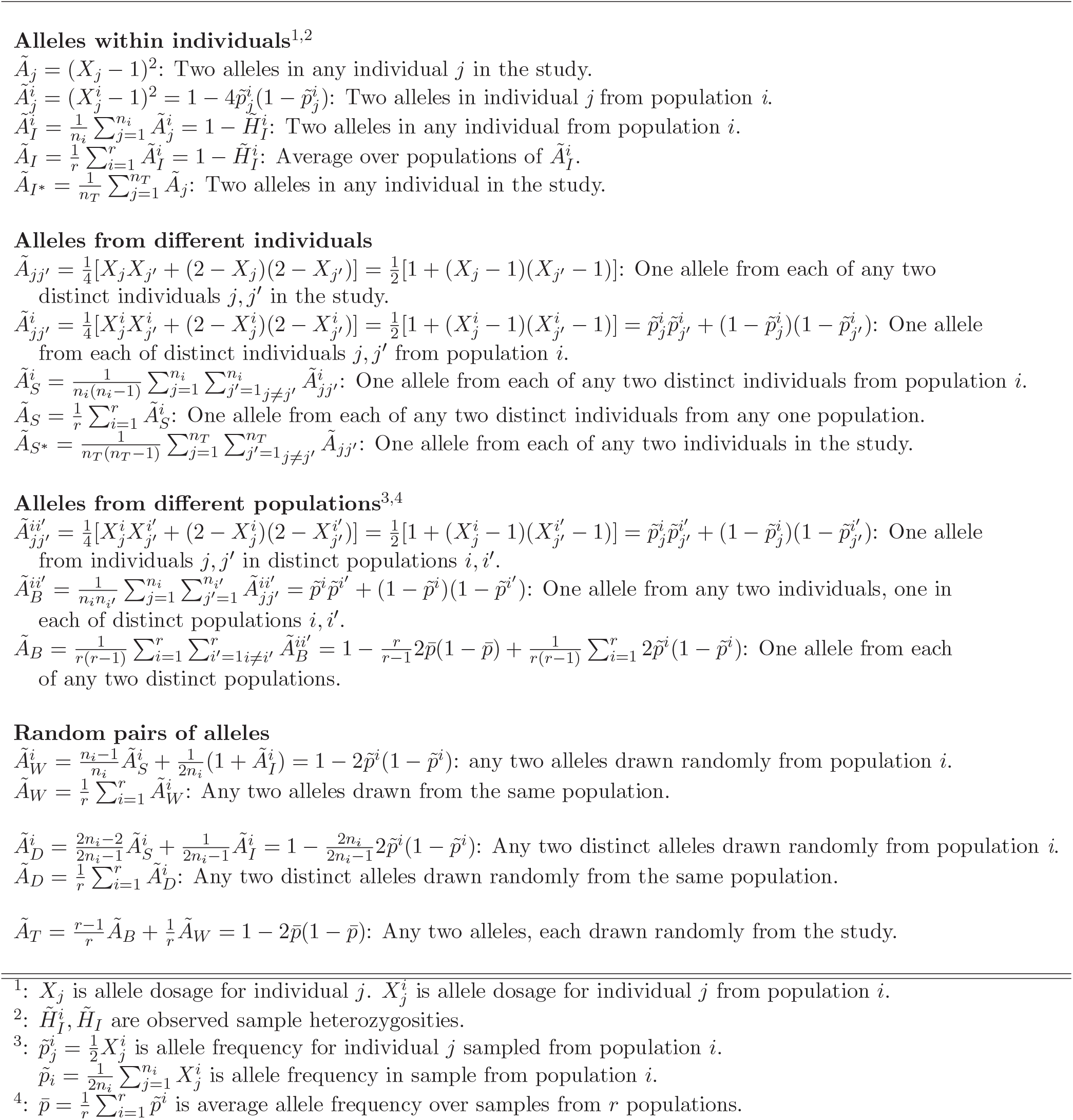

### Box 3: Allele-sharing Estimators (*Ã*’s from Box 2).

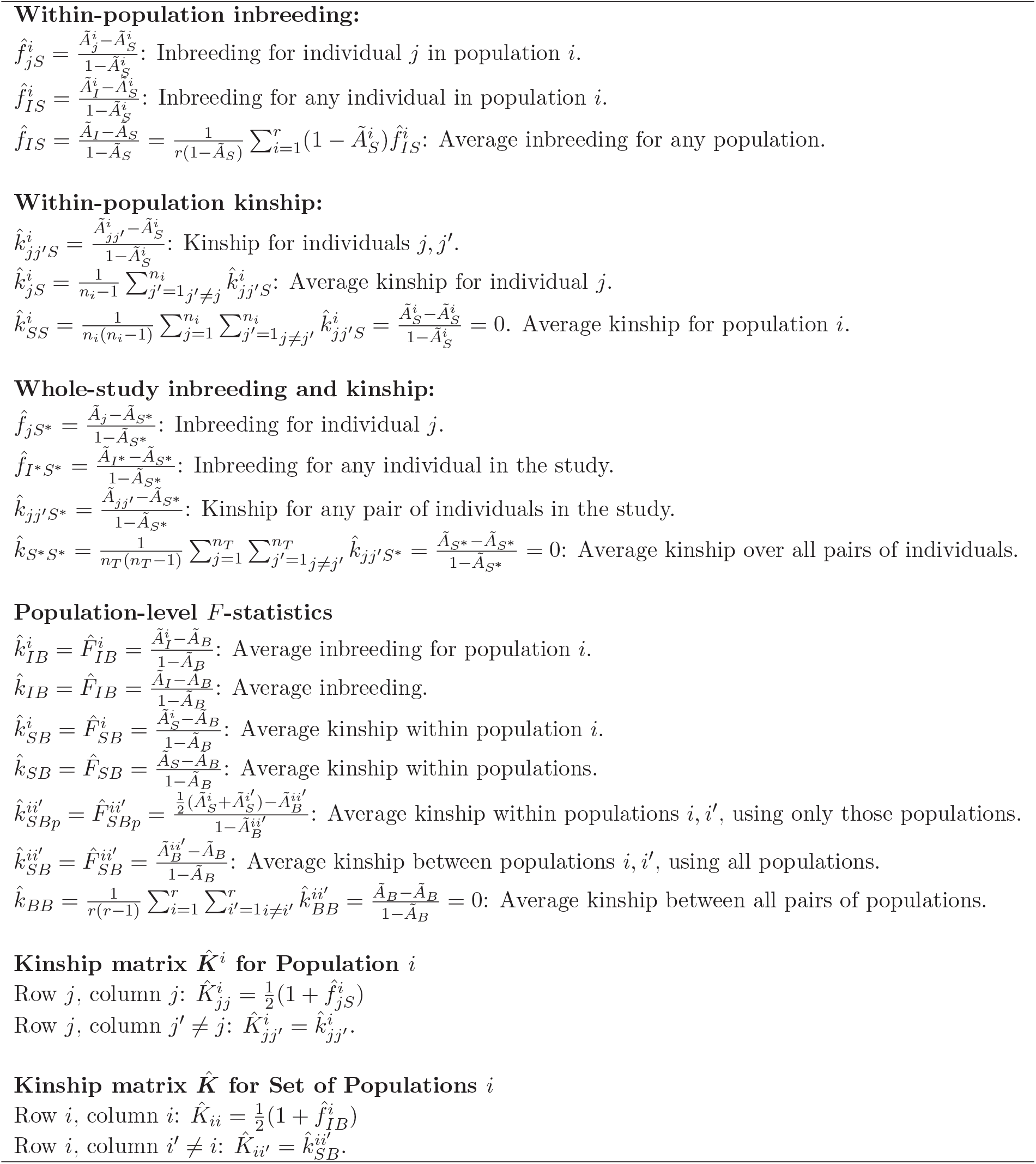

## References

Atag G, Waldman S, Carmi S, Soml M. 2024. An explanation for the sister replusion phenomenon in Patterson’s f-statistics. Genetics 228: iyae144 (2024). doi:10.1093/genetics/iyae144

Bhatia G, Patterson N, Sankararaman S, Price AK. 2013. Estimating and interpreting FST : The impact of rare variants. Genome Res. 23:1514–1521. doi:10.1101/gr.154831.113

Bilton TP. 2020. Developing Statistical Methods for Genetic Analysis of Genotypes from Genotyping-by-sequencing Data. Ph.D. Thesis, University of Otago. Available from https://hdl.handle.net/10523/9975.

Browning SR, Browning B. 2012. Identity by descent between distant relatives: Detection and applications. Ann Rev Genet. 46:617–633. doi:10.1038/541588-023-01582-w

Buckleton J, Curran J, Goudet J, Taylor D, Thiery A, Weir BS. 2016. Population-specific FST vales for forensic STR markers. Forens Sci Int: Genet. 23:91–100. doi:10.1016/j.fsigen.2016.03.004

Buckleton JS, Hall TO, Bright JA, Yung MC, Goudet J, Kruijver M, Weir BS. 2025. Estimation of population-specific values of theta for PowerPlex Y23 profiles. Forens Sci Int: Genet. 75: 103175 doi:10.1016/j.fsigen.2024.103175.

Byrska-Bishop M, Evani US, Zhao XF, Basile AO, Abel HJ, Regier AA, Corvelo A, Clarke WE, Musunuri R, Nagulapalli K, Fairley S, Runnels, A, Winterkorn L, Lowy E, Flicek P, Germer S, Brand H, Hall IM, Talkowski ME, Narzisi G, Zody MC. 2022. High-coverage whole-genome sequencing of the expanded 1000 Genomes Project cohort including 602 trios. Cell 185:3426–3440. doi 10.1016/j.cell.2022.08.004

Chesser RK. 1991. Gene diversity and female philopatry. Genetics 127:437–447. doi:10.1093/genetics/127.2.437

Cockerham CC. 1973. Analyses of gene frequencies. Genetics 74:679–700. doi:1093.genetics/74.4.679

Cockerham CC, Weir BS. 1987. Correlations, descent measures: Drift with migration and mutation. Proc Natl Acad Sci USA 84:8512–8514. doi:10.1073/pnas.84.23.8512

Conomos MP, Reiner AP, Weir BS, Thornton TA. 2016. Model-free estimation of recent genetic relatedness. Am J Hum Genet 98:127–148. doi:10.1016/j.ajhg.2015.11.022

Conrad DF, Jakobson M, Coop G, Wen X, Wall JD, Rosenberg MA, Prtichard JK. 2006. A worldwide survey of haplotype variation and linkage disequilibrium in the human genome. Nat Genet. 11:1251–1260. doi:10.1038/ng1911

Cotterman CW. 1940. Reprinted 1974. A calculus for statisticogenetics. Ph.D. thesis, Ohio State University, Columbus. In:Ballonoff P (ed) Genetics and Social Structure, Dowden: Hutchinson and Ross, Stroudsburg, Pennsylvania, pp 157–272.

Csűrös M. 2014. Non-identifiability of identity coefficients at biallelic loci. Theor Pop Biol. 92:22–29. doi:10.1016/j.tpb.2013.11.001

Cumer T, Machado AP, Siverio F, Cherkaoui SI, Roque I, Lourenço R, Charter M, Roulin A, Goudet J. 2022. Genomic basis of insularity and ecological divergence in barn owls (Tyto alba) of the Canary Islands. Heredity 129:281–294. doi:10.1038/s41437-022-00562-w

deBoer RA, Vega-Trejo R, Kotrschall A,Fitzpatrick JL. 2021. Meta-analytic evidence that animals rarely avoid inbreeding. Nat Ecol Evol. 5:949–964. doi:10.1038/s41559-021-01453-9

DeGiorgio M, Rosenberg NA. 2009. An unbiased estimator of gene diversity in samples containing related individuals. Mol Biol Evol. 26:501–512. doi:10.1093/molbev/msn254

Dowdeswell T. 2022. Forensic genetic genealogy: A profile of cases solved. Forens Sci Int :Genetics 58,102679. doi:10.1016/j.fsigen.2022.102679

Druet T, Gautier M. 2017. A model-based approach to characterize individual inbreeding at both global and local genomic scales. Mol Ecol. 26:5820–5841. doi:10.1111/mec.14324

Druet T, Gautier M. 2022. A hidden Markov model to estimate homozygous-by-descent probabilities associated with nested layers of ancestors. Theor Pop Biol. 145:38.41. doi:10.1016/j.tpb.2022.03.001

Evett IW. 1983. What is the probability that this blood came from that person: A meaningful question? J Forens Sci Soc. 23:35–59. doi:10.1016/S0015-7368(83)71540-9

Fan C, Mancuso N, Chiang CWK. 2022. A genealogical estimate of genetic relationships. Am J Hum Genet. 109:812–824. doi:101016/j.ajhg.2022.03.016

Gautier M, Vitalis R, Flori L, Estoup A. 2022. f-statistics estimation and admixture graph construction with Pool-Seq or allele count data using the R package poolfstat. Mol Ecol Resour. 22:1394–1416. doi:10.1111/1755-0998.13557

Gazal S, Sahbatou M, Perdry H, Letort S, Génin E, Leutenegger AL. 2014. Inbreeding coefficient estimation with dense SNP data: comparison of strategies and application to HapMap III. Hum Hered. 77:49–62. doi:10.1159/000358224

Gazal S, Sahbatou M, Babron MC, Génin E, Leutenegger AL. 2015. A high level of inbreeding in the final phase of 1000 Genome Project. Sci Rep. 5:17453. doi:10.1038/srep17453

Gibson J, Morton NE, Collins A. 2006. Extended tracts of homozygosity in outbred human populations. Hum Mol Genet. 15:789–795. doi:10.1093/hmg/ddi493

Goudet J. 2005. HIERFSTAT, a package for R to compute and test hierarchical F-statistics. Mol Ecol Notes 5:184–1896. doi:10.1111/j.1471-8286.2004.00828.x

Goudet J, Kay T, Weir BS. 2018. How to estimate kinship. Mol Ecol. 27:4121–4135. doi:10.1111/mec.14833

Goudet J, Weir BS. 2023. An allele-sharing, moment-based estimator of global, population-specific and population pair FST under a general model of population structure. PLoS Genet. 19:e01010871. doi:10.1371/journal.pgen.1010871

Graffelman J, Nelson S, Gogarten SM, Weir BS. 2015. Exact Inference for Hardy-Weinberg proportions with missing genotypes: Single and multiple imputation. G3 (Genes, Genomes, Genetics) 5:2365–2373. doi:10.1534/g3.115.022111

Graffelman J, Weir BS, Goudet J. 2025. Estimation of Jacquard’s genetic identity coefficients with bi-allelic variants by constrained least-squares. Heredity 134:10–20. doi:10.1038/s41437-024-00731-z

Guan Y, Levy D. 2024. Estimation of inbreeding and kinship coefficients via latent identity-by-descent states. BMC Bioinf. 40:btae082. doi:10.1093/bioinformatics/btae082

Gusev A. 2024. A molecular genetics perspective on the heritability of human behavior and group differences. Available for download at http://gusevlab.org/projects/hsq/hsq.pdf.

Harris K. 2023. Using enormous genealogies to map causal variants in space and time. Nat. Genet. 55:730–731. doi:10.1038/s41588-023-01389-9

Hartl DL, Clark AG. 2007. Principles of Population Genetics. 4th Edition. Sinuaer Associates, Sunderland MA.

Henn BM, Hn L, Macpherson JM, Eriksson N, Saxonov S, Pe’re I, Mountain JL. 2012. Cryptic distant relatives are common in both isolated and cosmopolitan genetic samples. PLoS ONE 7(4): e34267. doi:10.1371/journal.pone.0034267

Hill WG. 1996. Sewall Wright’s “Systems of Mating”. Genetics 143, 1499–1506. doi:10.1093/genetics/143.4.1499

Hill WG, Weir BS. 2011. Variation in actual relationship as a consequence of Mendelian sampling and linkage. Genet Res. 93:47–74. doi:10.1017/S0016672310000480

Hou Z, Ochoa A. 2023. Genetic association models are robust to common population kinship estimation biases. Genetics 224:iyad030. doi:10.1093/genetics/iyad030

Hudson RR, Slatkin M, Maddison WP. 1992. Estimation of levels of gene flow from DNA sequence data. Genetics 132:583–589. doi:10.1093/genefics/132.2.583

Jaquard A. 1972. Genetic information given by a relative. Biometrics 28:1101–1114. doi:10.2307/2528643

Jiang W, Zhou XY, Li ST, Song S, Zhao HY. 2022. An unbiased kinship estimation method for genetic data analysis. BMC Bioinf. 23:525–546. doi:10.1186/s12859-022-05082-2

Joshi PK, 2015. Directional dominance on stature and cognition in diverse human populations. Nature 523459–U176. doi:10.1038/nature14618

Künsch HR. 1989. The jackknife and bootstrap for general stationary observations. Ann Stat. 17:1217–1241. doi:10.1214/aos/1176347265

Lavanchy E, Goudet J. 2023. Effect of reduced genomic representation on using runs of homozygosity for inbreeding characterization. Mol Ecol Res. 23:787–802. doi:10.1111/1755-0998.13755

Lavanchy E, Weir BS, Goudet J. 2024. Detecting inbreeding depression in structured populations. Proc Natl Acad Sci USA 121:e2315780121. doi:10.1073/pnas.2315780121

Lehmann B, Lee H, Anderson-Trocmé L, Kelleher J, Gorjane G, Ralph PL. 2025. On ARGs, pedigrees, and genetic relatedness matrices. bioRxiv 2025.03 10.1011/2025.03.03.641310, preprint not peer reviewed.

Leslie S, Winney B, Hellenthal G, Davison D, Boumertit A, Day T, Hutnik K, Royrvik EC, Cunliffe B, Wellcome Trust Case Control Consortium, International Multiple Sclerosis Genitics Consortium, Lawson DJ, Falush D, Freeman C, Pirinen M, Myers S, Robinson M, Donnell P, Bodmer W. 2015. The fine-scale genetic structure of the British population. Nature 519:309–314. doi:10.1038/nature14230

Li CC, Horvitz DG. 1953. Some methods of estimating the inbreeding coefficient. Am J Hum Genet. 5:107–117. PMID:13065259; PMCID:PMC1716461

Malécot G. 1948. The Mathematics of Heredity. Translated by Yermanos DM. Freeman: San Francisco (1960).

Manichaikul A, Mychaleckyj JC, Rich SS, Daly K, Sale M, Chen WM. 2010. Robust relationship in genome-wide association studies. Bioinformatics 26:3867–2873. doi:10.1093/bioinformatics/btq559

Marsh WA, Brace S, Barnes I. 2023. Inferring biological kinship in ancient datasets: comparing the response of ancient DNA-specific software packages to low coverage data. BMC Genomics 24, 111 (2023). 10.1186/s12864-023-09198-4

Mary-Huard T, Balding D. 2023. Fast and accurate joint inference of coancestry parameters for populations and/or individuals. PLoS Genet. 10.1371/journal.pgen.1010054.

Meyermans R, Gorssen W, Buys N, Janssens S. 2020. How to study runs of homozygosity using PLINK. A guide for analyzing medium density SNP data in livestock and pet species. BMC Genomics 21,:4. doi:10.1186/s12864-020-6463-x

Milligan BG. 2003. Maximum-likelihood estimation of relatedness. Genetics 163:1153–1167. doi:10.1093/genetics/163.2.1153

Narasimhan V, Danecek P, Saclly A, Xue Y, Tyler-Smith C, Durbin R. 2016. BCFtools/RoH: a hidden Markov model approach for detecting autozygosity from next-generation sequencing data. Bioinformatics 32:1749–1752. doi:10.1093/bioinformatics/btw044

National Research Council. 2996. The Evaluation of Forensic DNA Evidence, National Academy Press, Washington DC.

Nei M. 1973. Analysis of gene diversity in subdivided populations. Proc Natl Acad Sci. USA 70:3321–3323. doi:10.1073/pnas.70.12.3321

Nei M. 1977. F-statistics and analysis of gene diversity in subdivided populations. Ann Hum Genet. 41:225–233. doi:10.1111/j.1469-1809.1977.tb01918.x

Nei M, Chesser RK. 1983. Estimation of fixation indices and gene diversities. Ann Hum Genet. 47:253–259. doi:10.1111/j.1469-1809.1983.tb00993.x

Nei M. 1986. Definition and estimation of fixation indices. Evolution 40:643–645. doi:10.2307/2408586

Nei M. 1987. Molecular Evolutionary Genetics. Columbia University Press, New York.

Nielsen R, Vaughn AN, Deng Y. 2025. Inference and applications of ancestral recombination graphs. Nat Rev Genet. 26:47–58. doi:10.1038/s41576-024

Ochoa A, Storey JD. 2021 Estimating FST and kinship for arbitrary population structures. PLoS Genet. 17:e1009241. doi:10.1371/journal.pgen.1009241

Ochoa A, Storey JD. 2023. Package ‘popkin’. https://cran.r-project.org/web/packages/popkin/popkin.pdf

Patterson N, Price AL, Reich D. 2006. Population structure and eigenanalysis. PLoS Genet. 2:e190. doi:10.1371/journal.pgen.0020190

Patterson N, Moorjani F, Luo Y, Mallik S, Rohland N, Zhan YP, Genschorek T, Webster T, Reich D. 2012. Ancient admixture in human history. Genetics 192:1065–1093. doi:10.1534/genetics.112.145037

Peter BM. 2016. Admixture, population structure and F-statistics. Genetics 202:1485–1501. doi:10.1534/genetics.115.183913

Reich A, Thangaraj K, Patterson N, Price AL, Singh L. 2009. Reconstructing Indian population history. Nature 461:489–494. doi:10.1038/nature08365

Reynolds J, Weir BS, Cockerham CC. 1983. Estimation of the coancestry coefficient: basis for a short term genetic distance. Genetics 105:767–779. doi:10.1093/genetics/105.3.767

Ringbauer H, Huang Y, Akbari A, Mallik S, Olalde I, Paterson N, Reich D. 2024. Nat Genet. 56:143–151. dor:10.1146/s41588-023-01582-w

Ritland K. 1996. Estimators for pairwise relatedness and individual inbreeding coefficients. Genet Res Camb. 67:175–185. doi:10.1017/S0016672300033620

Ritland K. 2024. Relatedness coefficients and their applications for triplets and quartets of genetic markers. G3 14:jkad236. doi:10.1093/g3journal/jkad236

Rousset F. 1996. Equilibrium values of measures of population subdivision for stepwise mutation models. Genetics 142:1357–1362. doi”10.1093/genetics/142.4/1357

Rousset F. 2002. Inbreeding and relatedness coefficients: what do they measure? Heredity 88:371–380. doi:10.1038/sj.hdy.6800065

Rousset F. 2007. Inferences from spatial population genetics. Chapter 28 in Balding DJ, Bishop M, Cannings C, (Editors). Handbook of Statistical Genetics 2,:945–979.

Ronfort J, Jenczewski E, Bataillon T, Roussett F. 1998. Ananlysis of population styructure in autotetrapolod species. Genetics 150:921–930. doi:10.1093/genetics/150.2.921

Ruitberg CM, Reeder DJ, Butler JM. 2001. STRBase: a short tandem repeat DNA database for the human identity testing community. Nucleic Acids Res. 29: 320–322. doi: 10.1093/nar/29.1.320

Shriver MD, Smith MW, Li J, Marcini A, Akey JM, Deka R, Ferrell RE. 1997. Ethnic-affiliation estimation by use of population-specific DNA markers. Am J Hum Genet. 60:957–64. PMCIDLPMC1712479

Speed D, Balding DJ. 2015. Relatedness in the post-genomic era: is it still useful? Nat Rev Genet. 16:33–44. doi:10.1038/nrg3821

The 1000 Genomes Project Consortium. 2010. A map of human genome variation from population-scale sequencing. Nature 467:1061–1073. doi:10.1038/nature09534

Thompson EA. 1975. The estimation of pairwise relationships. Ann Hum Genet. 39:173–189. doi:10.1111/j.1469-1809.1975.tb00120.x

Thompson EA. 2013. Identity by descent: variation in meiosis, across genomes, and in populations. Genetics 194:301–326. doi:10.1534/genetics.112.148825

Tsambos G. 2022. Efficient analyses of genetic ancestry in population-sized datasets. PhD thesis, The University of Melbourne. Accessible at https://hdl.handle.net/11343/311519

VanRaden, P.M. 2008. Efficient methods to compute genomic predictions. J Dairy Sci. 91:4414–4423. doi:10.3168/jds.2007-0980

Visscher PM, et al. 2006. Assumption-free estimation of heritability genome-wide identity-by-descent sharing between full siblings. PLoS Genetics 2:e41. doi: 10.1371/journal.pgen.0020041

Walsh B, Visscher P, Lynch M. 2025. Genetics and Analysis of Quantitative Traits. I Foundations, Oxford University Press.

Wang J. 2014. Marker-based estimates of relatedness and inbreeding coefficients: an assessment of current methods. J Evol Biol. 27:51–530. doi:10.1111/jeb.12315

Wang J. 2017. Estimating pairwise relatedness in a small sample of individuals. Heredity 119:302–313. doi:10.1038/hdy.2017.52

Wang J. 2022. A joint likelihood estimator of relatedness and allele frequencies from a small sample of individuals. Meth Ecol Evol. 13:2443–2462. doi:10.1111/2041-210X.13963

Wang J. 2025. EMIBD9: Estimating 9 condensed IBD coefficients, inbreeding and relatedness from marker genotypes. Heredity doi:10.1038/s41437-024-00739-5134:155–161.

Weir BS. 1996. Genetic Data Analysis II. Sinauer, Sunderland, MA.

Weir BS, Cardon LR, Anderson AD, Nielsen DM, Hill WG. 2005. Measures of human population structure show heterogeneity among genomic regions. Genome Research 15:1468–1476. doi10.1101/gr.4398405.

Weir BS, Cockerham CC. 1984. Estimating F-statistics for the analysis of population structure. Evolution 38:1358–1370. doi:10.1111/j.1558-5646.1984.tb05657.x.

Weir BS, Goudet J. 2017. A unified characterization of population structure and relatedness. Genetics 206:2085–2103. doi:10.1534/genetics.116.198424

Weir BS, Hill WG. 2002. Estimating F-statistics. Ann Rev Genet. 36:721–750. doi:10.1146/annurev.genet.36.050802.093940

Wong Y, Ignatieva A, Koskela J, Gorjanc G, Wohns AW, Kelleher J. 2024. A general and efficient representation of ancestral recombination graphs. Genetics 228:iyae100. doi:10.1093/genetics.iyae100

Wright S. 1922. Coefficients of inbreeding and relationship. Am Nat. 56:330–338.

Wright S. 1943. An analysis of local variability of flower color in Linanthus parryae. Genetics 28:139–156. doi:10.1093/genetics/28.2.139

Wright, S. 1951. The genetical structure of populations. Ann Eugenics 15:23–354. doi:10.1111/j.1469-1809.1949.tb02451.x

Wright, S. 1965. The interpretation of population structure by F-statistics with special regard to systems of mating. Evolution 19:395–420. doi:10.2307/2406450

Yang, J. Benyamin B McEvoy BH, Gordon S, Henders AK, Nyholt DR, Madden PA, Heath AC, Martin NG, Montgomery GW, Goddard ME, Visscher PM. 2010. Common SNPs explain a large proportion of the heritability for human height. Nat Genet. 42:565–569. doi:10.1038/ng.608

Yang J, Lee SH, Goddard ME, Visscher PM. 2011. GCTA: A tool for genome-wide complex trait analysis. Am J Hum Genet. 88:76–82. doi:10.1016/j.ajhg.2010.11.011

Yengo L, Zhu ZH, Wray NR Weir BS, Yang J, Robinson MR, Visscher PM. 2017. Detection and quantification of inbreeding depression for complex traits from SNP data. Proc Natl Acad Sci. USA 114:8602–8607. doi:10.1073/pnas.1621096114

Yu J, Pressoir G, Briggs WH, Bi IV, Yamasaki A, Doebley JF, McMullin MD, Gaut BS, Nielsen DM. Holland JB, Kresovich S, Buckler ES. 2006. A unified mixed-model method for association mapping that accounts for multiple levels of relatedness. Nat Genet. 38:203–207. doi:10.1038/ng1702

Zhang BC, Biddanda A, Gunnarsson AF, Cooper F, Palamara PF. 2023. Biobank-scale inference on ancestral recombination graphs enables genealogical analysis of complex traits. Nat. Genet. 55:768–776. doi:10.1038/s41588-023-01379-x

Zhang Q, Goudet J, Weir BS. 2022. Rank-invariant estimation of inbreeding coefficients. Heredity 128:1–10. doi:10.1038/s41437-021-00471-4

